# Modulation of the Cardiac Sodium Channel Na_V_1.5 Peak and Late Currents by NAD^+^ Precursors

**DOI:** 10.1101/2020.02.20.958066

**Authors:** Daniel S. Matasic, Jin-Young Yoon, Jared M. McLendon, Haider Mehdi, Mark S. Schmidt, Alexander M. Greiner, Pravda Quinones, Gina M. Morgan, Ryan L. Boudreau, Kaikobad Irani, Charles Brenner, Barry London

## Abstract

**Rationale:** The cardiac sodium channel Na_V_1.5, encoded by *SCN5A*, produces the rapidly inactivating depolarizing current I_Na_ that is responsible for the initiation and propagation of the cardiac action potential. Acquired and inherited dysfunction of Na_V_1.5 results in either decreased peak I_Na_ or increased residual late I_Na_ (I_Na,L_), leading to tachy/bradyarrhythmias and sudden cardiac death. Previous studies have shown that increased cellular NAD^+^ and NAD^+^/NADH ratio increase I_Na_ through suppression of mitochondrial reactive oxygen species and PKC-mediated Na_V_1.5 phosphorylation. In addition, NAD^+^-dependent deacetylation of Na_V_1.5 at K1479 by Sirtuin 1 increases Na_V_1.5 membrane trafficking and I_Na_. The role of NAD^+^ precursors in modulating I_Na_ remains unknown.

**Objective:** To determine whether and by which mechanisms the NAD^+^ precursors nicotinamide riboside (NR) and nicotinamide (NAM) affect peak I_Na_ and I_Na,L_ *in vitro* and cardiac electrophysiology *in vivo*.

**Methods and Results:** The effects of NAD^+^ precursors on the NAD^+^ metabolome and electrophysiology were studied using HEK293 cells expressing wild-type and mutant Na_V_1.5, rat neonatal cardiomyocytes (RNCMs), and mice. NR increased I_Na_ in HEK293 cells expressing Na_V_1.5 (500 μM: 51 ± 18%, p=0.02, 5 mM: 59 ± 22%, p=0.03) and RNCMs (500 µM: 60 ± 26%, p=0.02, 5 mM: 75 ± 39%, p=0.03) while reducing I_Na,L_ at the higher concentration (RNCMs, 5 mM: −45 ± 11%, p=0.04). NR (5 mM) decreased Na_V_1.5 K1479 acetylation but increased I_Na_ in HEK293 cells expressing a mutant form of Na_V_1.5 with disruption of the acetylation site (Na_V_1.5-K1479A). Disruption of the PKC phosphorylation site abolished the effect of NR on I_Na_. Furthermore, NAM (5 mM) had no effect on I_Na_ in RNCMs or in HEK293 cells expressing wild-type Na_V_1.5, but increased I_Na_ in HEK293 cells expressing Na_V_1.5-K1479A. Dietary supplementation with NR for 10-12 weeks decreased QTc in C57BL/6J mice (0.35% NR: −4.9 ± 2.0%, p=0.26; 1.0% NR: −9.5 ± 2.8%, p=0.01).

**Conclusions:** NAD^+^ precursors differentially regulate Na_V_1.5 via multiple mechanisms. NR increases I_Na_, decreases I_Na,L_, and warrants further investigation as a potential therapy for arrhythmic disorders caused by Na_V_1.5 deficiency and/or dysfunction.

## 1. INTRODUCTION

The main cardiac sodium channel Na_V_1.5 generates a rapid inward depolarizing Na^+^ current (I_Na_) that initiates and allows propagation of the cardiac action potential throughout the heart.^1^ Loss-of-function mutations in *SCN5A*, the gene encoding the pore-forming alpha subunit of Na_V_1.5, result in: (1) a reduction of I_Na_, (2) a decrease in conduction velocity, and (3) an increased risk for tachyarrythmias, bradyarrhythmias, and sudden cardiac death.^2–5^ Inherited deficiency in Na_V_1.5 can manifest as several conditions including Brugada Syndrome, sick sinus syndrome, and progressive cardiac conduction defects, whereas gain-of-function mutations in *SCN5A* that limit voltage- and time-dependent inactivation increase late Na^+^ current (I_Na,L_) and result in Long QT syndrome type 3 (LQT3).^6–9^ Moreover, acquired dysfunction of Na_V_1.5 in conditions such as cardiomyopathies and heart failure can decrease I_Na_ and increase I_Na,L_, contributing to fatal arrhythmias.^10^ Given the importance of Na_V_1.5 in modulating arrhythmic risk, efforts are underway to modulate Na_V_1.5 expression and/or activity as a potential therapeutic avenue for arrhythmogenesis in inherited and acquired conditions.

The cellular redox coenzyme nicotinamide adenine dinucleotide (NAD^+^) has been established as an important regulator of cardiac physiology and pathophysiology.^11^ Previous studies have shown that NAD^+^ supplementation can normalize the NAD^+^/NADH redox ratio and increase I_Na_.^12^ Additionally, NAD^+^ supplementation and increased NAD^+^/NADH ratio have been shown to alter protein kinase A (PKA) and protein kinase C (PKC) dependent Na_V_1.5 phosphorylation, thereby altering Na_V_1.5 surface expression and/or channel conductance.^12–14^ More recently, we showed that the NAD^+^-dependent deacetylase Sirtuin-1 (SIRT1) increases Na_V_1.5 surface expression and I_Na_ by deacetylating a lysine residue (K1479) within the Na_V_1.5 III-IV intracellular linking domain *in vitro* and in animal models.^15, 16^

Given that NAD^+^ has poor bioavailability, NAD^+^ precursors are emerging as a therapeutic strategy for the prevention of cardiovascular disease.^11^ Nicotinamide Riboside (NR), a bioavailable and well-tolerated oral NAD^+^ precursor, has emerged as a leading NAD^+^ supplement in humans. Several ongoing clinical trials examining the effects of NR supplementation on cardiovascular diseases including heart failure are currently underway.^17–20^ However, the effects of NAD^+^ precursor supplementation on Na_V_1.5 function and cardiac electrophysiology are relatively unexplored.

In this report, we demonstrate that the NAD^+^ precursor NR alters the NAD^+^ metabolome, increasing NAD^+^ content, Na_V_1.5 deacetylation, and peak I_Na_ while reducing I_Na,L_. In addition, nicotinamide (NAM), an alternative NAD^+^ precursor known to inhibit sirtuin activity at high doses, does not increase I_Na_ at high dose in either HEK293 cells or rat neonatal cardiomyocytes (RNCMs). We demonstrate that the effect of NR is mediated by the PKC-phosphorylation S1503 site on Na_V_1.5 and that the inhibitory effect of NAM is mediated by the K1479 acetylation site. Furthermore, dietary supplementation of 1% NR in wild-type mice resulted in a reduction in QTc after 10-12 weeks, consistent with the reduction in I_Na,L_. Together, this work lays a foundation for further investigation of the NAD^+^ precursor NR as a potential therapeutic agent for the prevention of arrhythmias in cardiac diseases associated with Na_V_1.5 deficiency and/or dysfunction.

## 2. METHODS

All animal studies were approved by the Institutional Animal Care and Use Committees (IACUC) at the University of Iowa.

### 2.1 Cell Culturing of Non-Myocytes and Myocytes

HEK293 cells and HEK293 cells stably-expressing Na_V_1.5 were obtained from the American Type Culture Collection (ATCC, Manassas, VA) and Dr. Samuel Dudley (University of Minnesota), respectively, and cultured in 10%-fetal bovine serum and 1% penicillin/streptomycin supplemented Dulbecco’s Modified Eagle’s Medium (DMEM). RNCMs were isolated from 1-3 day old Sprague-Dawley rat pups using a neonatal cardiomyocyte isolation system (Worthington Biochemical Co, Lakewood, NJ) and cultured in 5%-horse serum supplemented DMEM/F-12 equivolumetric mixture.

### 2.2 Reagents

Transient expression of Na_V_1.5 constructs in HEK293 cells was achieved utilizing Lipofectamine 2000 (Invitrogen, Carlsbad, CA). Mutant Na_V_1.5-K1479A and Na_V_1.5-K1479A/S1503A were generated by site-directed mutagenesis using the Quik-Change II XL kit (Agilent Technologies, Santa Clara, CA). The following NAD^+^ metabolites were utilized: nicotinamide riboside (NR, Niagen, Irvine, CA), nicotinamide (NAM, Sigma-Aldrich, St. Louis, MO), and 1-methlynicotinamide (meNAM, Sigma-Aldrich, St. Louis, MO). In experiments using transient transfection of Na_V_1.5, treatment of NAD^+^ metabolites was initiated 12 hours post-transfection, continued for 48 hours, and cells were subjected to whole-cell patch clamp after plating onto poly-D-lysine-coated coverslips. In experiments using RNCMs, treatment with NAD^+^ metabolites was continued for 24 hours post-plating onto laminin-coated coverslips.

### 2.3 Whole-Cell-Patch Clamp of Na^+^ Currents (I_Na_)

All whole-cell recordings were obtained utilizing the Axon Axopatch 200B amplifier and Digidata 1440B data acquisition system (Molecular Devices, San Jose, CA), as previously described.^15^ Treatment with NAD^+^ metabolites was continued throughout experimentation with extracellular solutions containing the respective concentration of the respective metabolite. Cell capacitance was recorded directly from the amplifier after adjusting for the transient post-membrane rupture.

For HEK293 cells, patch pipettes of 2-3 MΩ were filled with an internal solution containing: 110 mM CsF, 20 mM CsCl, 10 mM NaF, 10 mM EGTA, 10 mM HEPES, with the pH adjusted to 7.35 with CsOH. The extracellular solution contained: 25 mM NaCl, 128 mM NMDG, 4.5 mM KCl, 10 mM HEPES, 1 mM MgCl_2_, 1.5 mM CaCl_2_, 5 mM glucose, and pH adjusted to 7.35 using HCl. To test the steady-state activation of I_Na_ in HEK293 cells, a 500-ms prepulse to −120 mV was initiated to recover any channels from inactivation and then cells were subjected to a 20 or 200-ms test pulse between −90 mV and +40 mV in increments of 5 mV. The peak current density was assessed at the test potential of largest inward current. To test the steady-state inactivation, a two-pulse protocol was employed with 500-ms conditioning pulses varying from −140 mV to −30 mV, followed by a 20-ms test pulse at −20 mV.

For RNCMs, patch pipettes of 2-3 MΩ were filled with an internal solution containing: 10 mM NaCl, 90 mM aspartic acid, 70 mM CsOH, 10 mM EGTA, 20 mM CsCl, 10 mM HEPES, and pH adjusted to 7.35 using CsOH. The extracellular solution contained: 25 mM NaCl, 120 mM CsCl, 4.5 mM KCl, 10 mM HEPES, 2 mM MgCl_2_, 0.5 mM CaCl_2_, and pH adjusted to 7.35 using CsCl. To test the steady-state activation of I_Na_ in RNCMs, a 200-ms prepulse to −120 mV was used to eliminate any inactivated channels and then cells were subjected to a 200-ms test pulse between −80 mV and +15 mV in increments of 5 mV. To test the steady-state inactivation, a two-pulse protocol was employed with 1-s conditioning pulses varying from −150 mV to +30 mV followed by a 200-ms test pulse at −20 mV. I_Na,L_ was calculated by averaging the current between 50-ms and 150-ms of the 200-ms depolarization at three test-potentials: 20 mV, 25 mV, 30 mV). To verify the measurements of I_Na,L_ in RNCMs, a 143 mM NaCl extracellular solution was used and RNCMs were subjected to a continuous protocol of a 200-ms prepulse to −120 mV followed by a 200-ms depolarizing test-pulse to −20 mV with a 5-s interpulse duration at a holding potential of −80 mV before and after application of 10 µM tetrodotoxin (TTX). TTX-sensitive I_Na,L_ was calculated by the difference of measured late current before and after TTX.

pClamp software (version 10.4) was utilized for data analysis. To calculate I_Na_, peak current was normalized to the membrane capacitance. Steady-state activation and inactivation curves were generated and fitted to the Boltzmann equation, whereupon half-maximal potential (V_1/2_) and slope factor (*k*) were derived.

### 2.4 Whole-Cell-Patch Clamp of K^+^ and Ca^2+^ Currents

Conventional whole-cell patch clamp techniques were used to record K^+^ and Ca^2+^ currents from RNCM. Total potassium current was evoked during 4-s depolarizing voltage steps to potentials between −110 and 50 mV from a holding potential of −80 mV in 10 mV increments. Each trial was preceded by a short (20-ms) depolarization to −20 mV to eliminate contamination of Na^+^ currents. I_K,total_ was defined as the currents taken 20-ms into depolarization, and I_K,sustained_ as the currents taken just prior to repolarization. I_to_ was determined by the difference between I_K,total_ and I_K,sustained_. Rapidly activating, slowly inactivating outward current (I_Kur_) was obtained by subtraction of currents evoked by depolarizing voltage steps (−110 to 50 mV in 10 mV increments) from a holding potential of −40 mV and −80 mV followed by a 100-ms −40 mV inactivating prepulse. Ca^2+^ current (I_Ca_) was elicited by holding at −80 mV and stepping between −70 and 70 mV (10 mV steps) for 500-ms after a prepulse to −40 mV for 1-s to inactivate Na^+^ currents.

The internal pipette solution for I_K_ voltage clamp studies was: 140 mM KCl, 4 mM Mg-ATP, 1 mM MgCl_2_, 5 mM EGTA, 10 mM HEPES; the extracellular solution contained: 143 mM NaCl, 4.5 mM KCl, 10 mM HEPES, 2.5 mM CaCl_2_, 1 mM MgCl_2_, and 0.1 mM CdCl_2_. The pH was adjusted to 7.4 with NaOH. The internal pipette solution for I_Ca_ recording was: 110 mM CsCl, 20 mM TEACl, 5 mM Mg-ATP, 2 mM Na_2_-ATP, 5 mM Na Creatine Phosphate, 5 mM HEPES, 3 mM EGTA, and pH was adjusted to 7.2 with TRIS. The extracellular solution was: 130 mM NMDG, 20 mM Tetraethylammonium chloride (TEACl), 2 mM CaCl_2_, 2 mM MgCl_2_, 10 mM HEPES, 10 mM Glucose, and the pH was adjusted to 7.4 with HCl. Cells were dialyzed for 5 min prior to initiating experimental protocols.

### 2.5 Quantification of the NAD^+^/NADH Ratio and the NAD^+^ Metabolome

NAD^+^, NADH, and NAD^+^/NADH ratios was quantified in cellular extracts using the NAD/NADH-Glo™ Assay Kit (Promega Co, Madison, WI). Direct measurements of NAD^+^ and other metabolites in the NAD^+^ metabolome were performed on cell extracts using Liquid Chromatography-Mass Spectrometry (LC-MS), as previously described.^21^ Measurements below the quantitation limit were set to the minimum value of quantitation.

### 2.6 Western Blotting

Conventional techniques were utilized as previously described.^15^ To assess the acetylation of K1479, a custom-designed anti-acetyl K1479 Na_V_1.5 antibody was utilized, as previously described and used.^15^ Total Na_V_1.5 expression in whole cell lysates (Alomone ASC-005) was normalized to GAPDH (<u>Trevigen</u> 2275-PC). To measure the membrane expression of Na_V_1.5, the membrane-enriched protein fraction was harvested with the Compartmentalization Protein Extraction Kit (Millipore, Burlington, MA).

### 2.7 Assessment of Na_V_1.5 Surface Expression by Microscopy

HEK293 cells that stably express Na_V_1.5 with an extracellular FLAG epitope in the extracellular region of Domain I were seeded on gelatin coated glass bottom 24-well plates (Cellvis, Sunnyvale, CA). These FLAG-Na_V_1.5 HEK293 cells were treated for 48 hours with 5 mM NR (replenished media at treatment at 24 hours) and subsequently fixed for 15 minutes with 4% paraformaldehyde (diluted from 16%, Electron Microscopy Sciences, Hatfield, PA). Without cell permeabilization, cells were incubated in blocking buffer (PBS with 2% BSA 5% goat serum) for 1 hour, and then incubated overnight with Anti-FLAG M2 Antibody (Sigma Aldrich, St. Louis MO), diluted 1:1000 in blocking buffer, followed by a one hour incubation in Alexa-568 conjugated anti-mouse antibody (Abcam, Cambridge UK) diluted 1:1000 in blocking buffer. Surface Expression of FLAG-Nav1.5 was assessed by confocal microscopy using a Zeiss LSM510 microscope with a 63x objective (Plan Apochromat 63x/1.4 oil DIC) and Zeiss ZEN software.

### 2.8 Dietary Supplementation of Nicotinamide Riboside in Mice

4-5-month-old C57BL/6J mice of both sexes were randomly placed on a control diet (Teklad 2920X, Envigo) or a diet supplemented with NR at either 0.35% or 1% (3.5 g or 10 g of NR, respectively, in 1 kg of Teklad 2920X, Envigo) for 10-12 weeks. Food was prepared fresh every 3-4 days and mice were weighed to assess for any changes in body weight throughout the experiment.

### 2.9 Electrocardiography and Echocardiography in Mice

Electrocardiograms (EKGs) were performed at baseline, 4-6 weeks of diet, and 10-12 weeks of diet using the iWork IX/100B EKG recording system. The mice were anesthetized with 1-2% isoflurane, and high resolution multi-lead EKGs were obtained by placing bipolar electrodes subcutaneously in positions corresponding to human leads: I (right to left arm), II (right arm to left leg), III (left arm to left leg), and modified chest V (back to left sternal border), as previously described.^22^ Data was collected, digitized, and analyzed using LabScribe 2.0 software (iWorx, Dover, NH). An investigator blinded by treatment group analyzed R-R interval, PR interval, QRS duration, and QT interval for leads I, II, III, and V. QTc was calculated as previously described.^23^ Maximum values among leads for PR, QRS, and QTc are reported.

Transthoracic echocardiography was performed to assess cardiac size and function at baseline and 10-12 weeks of diet (Vevo 2700 VisualSonics System). Mice were anesthetized with midazolam; measurements included heart rate, end diastolic volume, end systolic volume, and ejection fraction as previously described.^24^

### 2.10 Statistical Analysis

All statistical analysis was performed with GraphPad Prism. Results are represented as the mean ± standard error of the mean and considered statistically significant if p-values were ≤ 0.05. All analyses were performed blinded to treatment or construct, as appropriate. Significance of difference between two groups was evaluated using independent sample t-tests. In addition, two-way ANOVAs were used to evaluate differences between groups over time.

## 3. RESULTS

### 3.1 Nicotinamide Riboside Increases I_Na_ in HEK293 Cells Expressing Na_V_1.5 and in RNCMs

Previous studies showed that extracellular application of NAD^+^ reversed the downregulation of I_Na_ that resulted from increased NADH levels in HEK293 cells expressing Na_V_1.5.^12^ We tested whether NR (Figure 1A), a bioavailable precursor of NAD^+^, can influence I_Na_. NR (500 μM) supplementation of HEK293 cells constitutively expressing Na_V_1.5 increased NAD^+^ and NADH levels without changing the NAD^+^/NADH ratio (Figure 1B), and increased I_Na_ (51 ± 18%, p=0.02; Figure 1C-E, Table S1) with only minor changes to Na_V_1.5 gating properties (4 mV hyperpolarizing shift in the steady-state activation half-potential; Figures 1F, S1). Similarly, 5 mM NR significantly increased I_Na_ in HEK293 cells transiently-transfected with the wild-type Na_V_1.5 channel (59 ± 22%, p=0.03, Figure 1G-H, Table S1) with no effects on their gating properties (Figures 1I, S1, Table S5). Surface expression of the channel was not increased in response to NR (Figures S2, S3).

**Figure 1.**
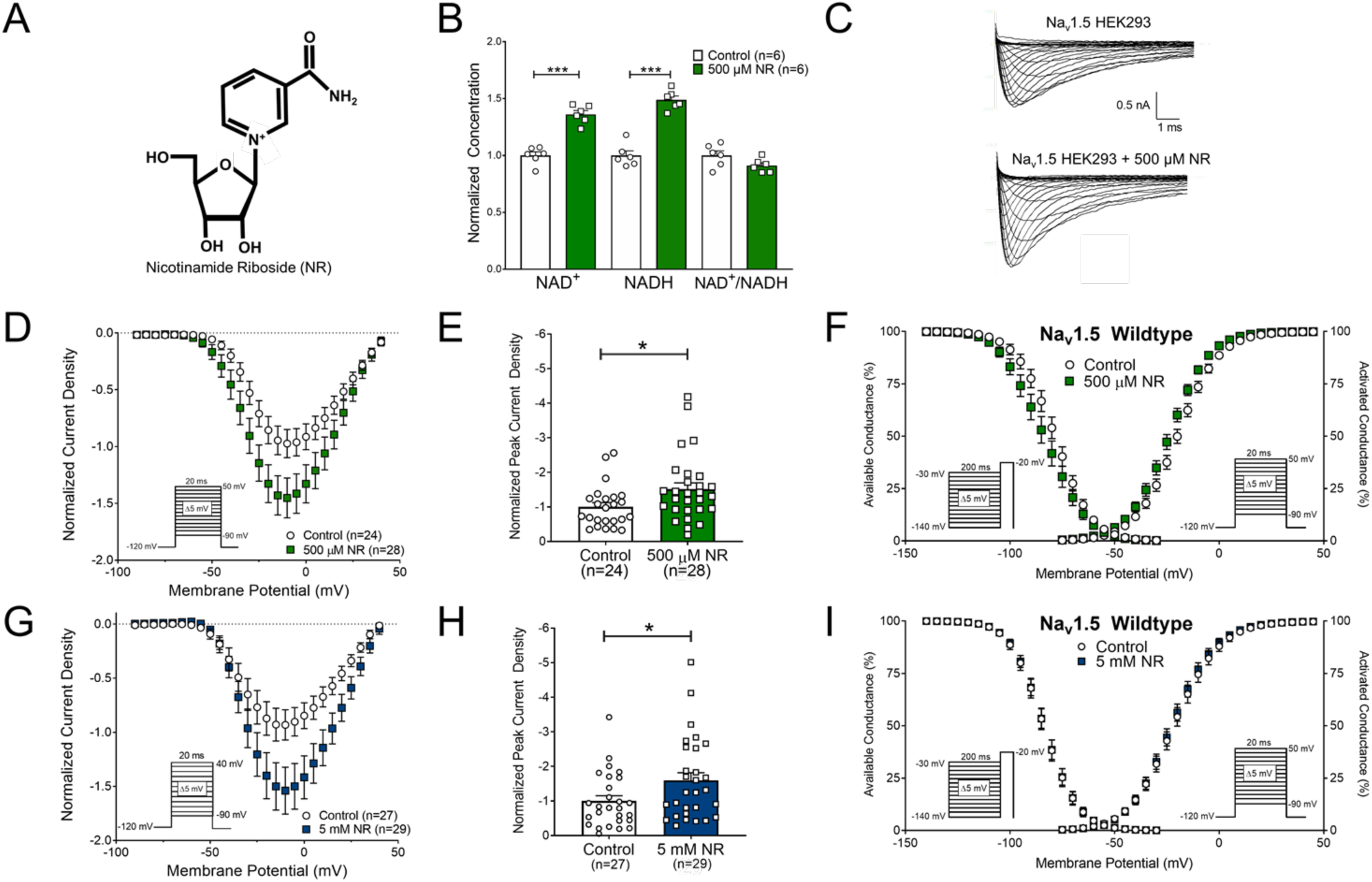
Nicotinamide Riboside (NR) increases I_Na_ without a change in NAD^+^/NADH ratio in HEK293 cells stably-expressing and transiently-expressing Na_V_1.5. **(A)** Structure of NR, **(B)** Quantification of NAD^+^ and NADH demonstrating significantly elevated levels of both metabolites, without altering the NAD^+^/NADH ratio, with NR supplementation **(C)** Representative traces of I_Na_, **(D)** current-voltage (I-V) relationship, and **(E)** normalized peak current density between HEK293 cells stably-expressing Na_V_1.5 with and without NR supplementation (500 µM, 48 hours) illustrate that NR increases I_Na_. **(F)** Gating properties including steady-state activation and inactivation are relatively unchanged, with only a 4 mV hyperpolarizing shift in activation, with NR supplementation. **(G-I)** Similar results are observed in HEK293 cells transiently-transfected with Na_V_1.5 (5 mM NR, 48 hours). Differences between groups were statistically determined utilizing unpaired t-tests (*p<0.05, ***p<0.001).

Comparable results were observed in RNCMs (Figure 2). Representative whole-cell traces and I-V curves illustrated the stimulatory effects of NR on I_Na_ (500 µM NR: 60 ± 26%, p=0.02; 5 mM NR: 74 ± 39%, p=0.03; Figure 2A-C, Table S2). Changes in gating properties including steady-state activation and inactivation were observed after treatment with 5 mM NR but not 500 µM NR, with a depolarizing shift of steady-state activation and inactivation (Figures 2D, S4, Table S5). Interestingly, 5 mM NR reduced I_Na,L_ (−45 ± 11%, p=0.04, measured at −25 mV; Figure 2E-F). The absolute late current densities, measured at −25 mV, for 5 mM NR relative to control (−1.88 ± 0.23 pA/pF vs. −3.32 ± 0.26 pA/pF, p=0.002, Figure S5) also decreased. To verify that the observed changes in I_Na,L_ were not attributed to contaminated leak currents, I_Na,L_ was also determined by measuring the TTX-sensitive current. 5 mM NR significantly decreased the TTX-sensitive I_Na,L_ compared to control in RNCMs (−1.98 ± 0.48 pA/pF vs. −4.43 ± 1.24 pA/pF, p=0.04, Figure 2G-H). NR increased NAD^+^ and NADH content in RNCMs, similar to the findings in HEK293 cells; however, NR also significantly increased the NAD^+^/NADH ratio in RNCMs (Figure 2I).

**Figure 2.**
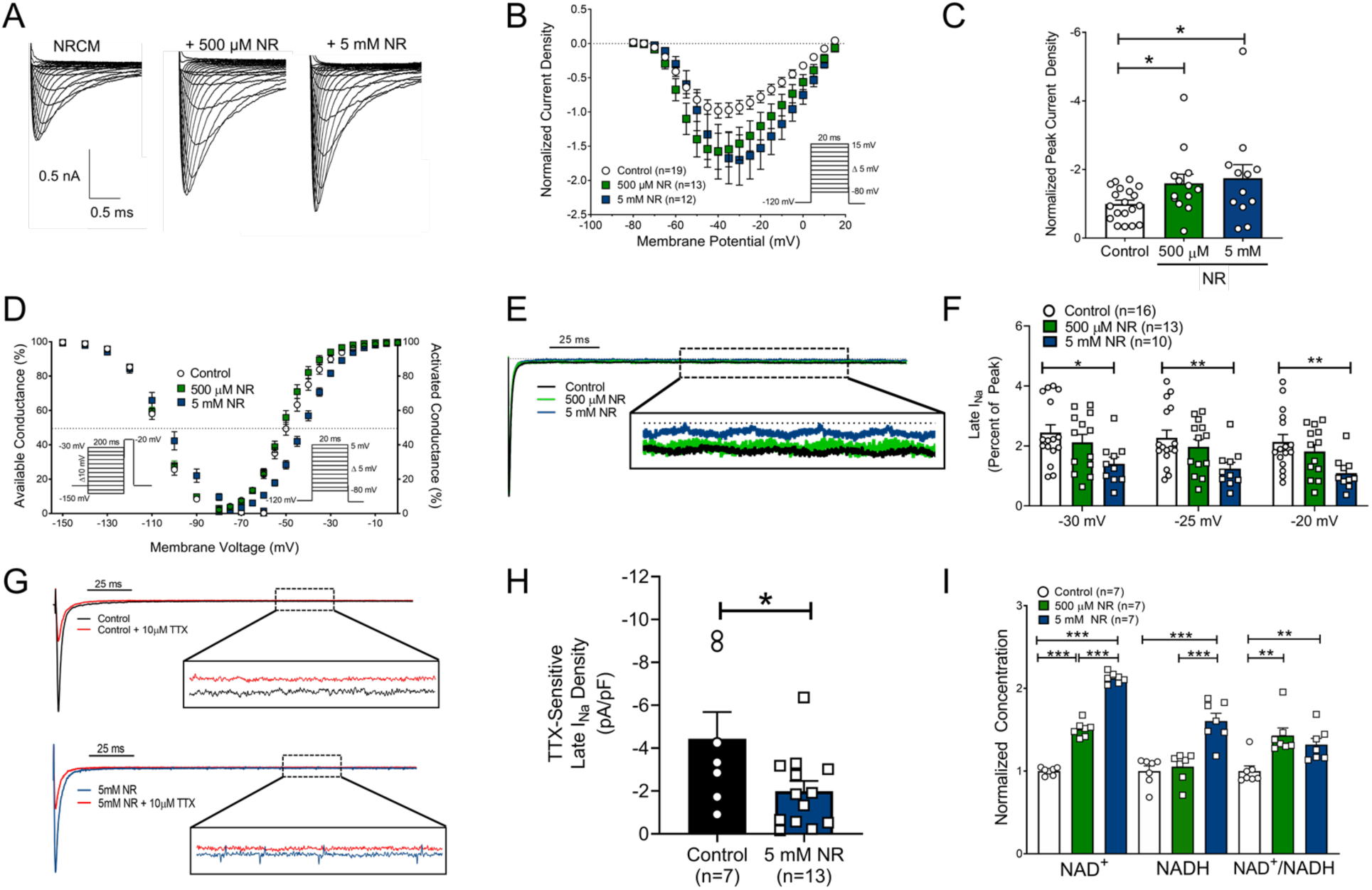
Nicotinamide Riboside (NR) increases I_Na_ with dose-dependent effects on gating properties in Neonatal Rat Cardiomyocytes. **(A)** Representative current traces, **(B)** I-V curve, and **(C)** normalized peak current density of rat myocytes treated with NR for 48 hours. **(D)** Gating properties of steady-state activation and inactivation **(E-F)** Representative traces and quantification of I_Na,L_ **(G-I)** Representative traces and quantification of tetrodotoxin (TTX)-sensitive changes in I_Na,L_. **(I)** NAD^+^, NADH, and NAD^+^/NADH ratio quantified in rat myocytes after 24-hour NR treatment (500 µM or 5 mM). Differences between groups were statistically determined utilizing unpaired t-tests (*p<0.05, **p<0.01, ***p<0.001).

### 3.2 Nicotinamide Riboside Drives Deacetylation of Na_V_1.5

Our group recently showed that the NAD^+^-consuming enzyme Sirtuin 1 regulates Na_V_1.5 trafficking by deacetylation of a lysine residue at the 1479 position within the III-IV intracellular linker domain.^15^ NR supplementation of HEK293 cells transiently transfected with Na_V_1.5 decreased the acetylation of K1479 only at the higher 5 mM dose, with no effect on total Na_V_1.5 mRNA or protein expression (Figures 3, S6).

**Figure 3.**
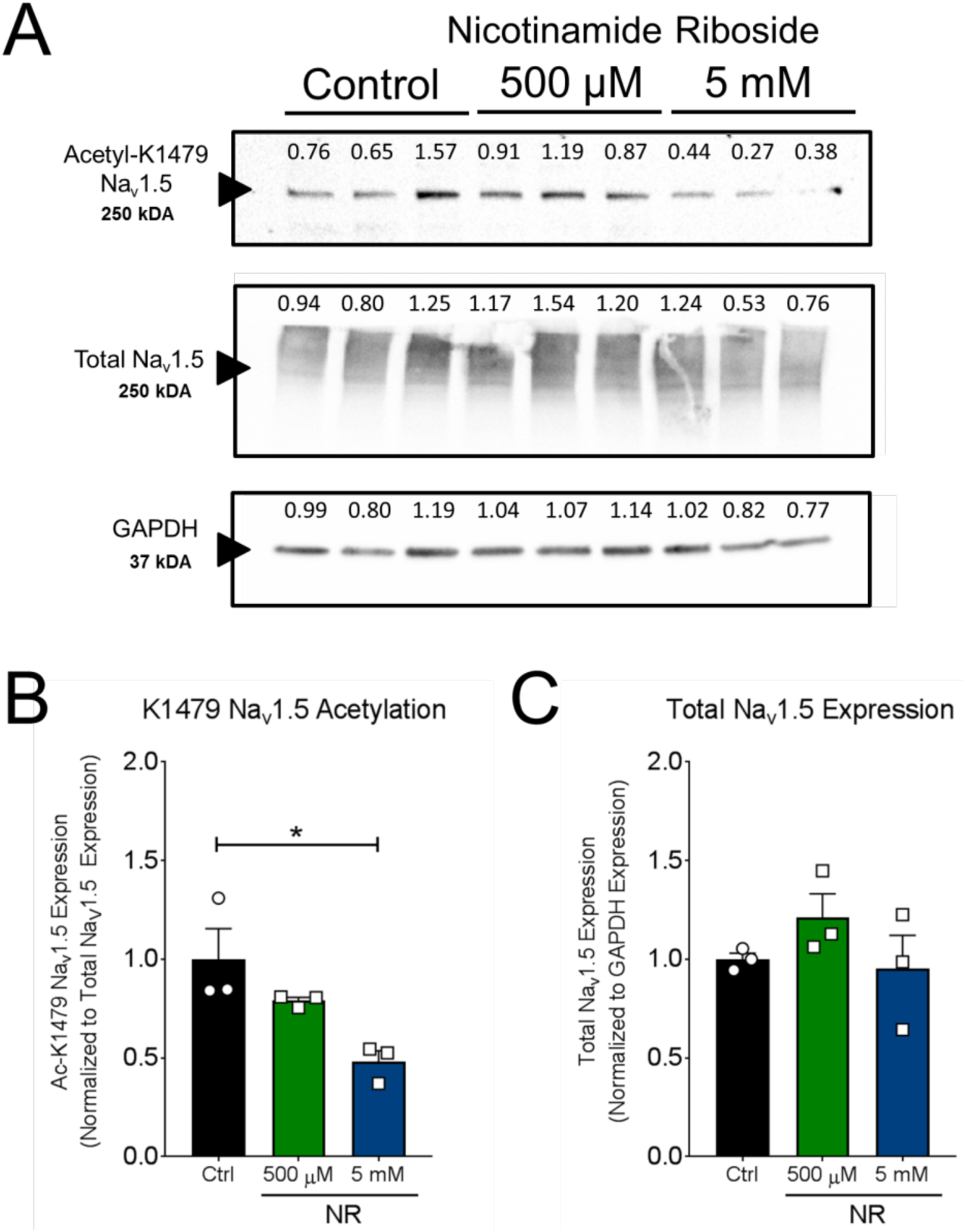
Nicotinamide Riboside supplementation decreases K1479 acetylation in Na_V_1.5-HEK293 cells. **(A)** Acetylated K1479 Na_V_1.5, total Na_V_1.5, and GAPDH expression in HEK293 cells transiently-transfected cells Na_V_1.5 and treated with NR (500 µM or 5 mM, 48 hours) or H_2_O. **(B)** Acetylated K1479 was normalized to total Na_V_1.5 expression and **(C)** total Na_V_1.5 expression was normalized to GAPDH expression. Differences between groups were statistically determined utilizing One-Way ANOVA tests with multiple comparisons (*p<0.05).

### 3.3 Nicotinamide Riboside does not alter K^+^ currents or Ca^2+^ currents

To assess whether NR has an influence on other currents regulating the cardiac action potential, K^+^ currents and Ca^2+^ currents were assessed by whole-cell patch clamp in RNCMs after a 24 hour treatment with 5 mM NR. NR had no significant effect on K^+^ currents including total I_K_, sustained I_K_, I_to_, and I_Kur_. NR had no effect on Ca^2+^ current density and inactivation kinetics in RNCMs as assessed by whole cell patch clamp (Figure S7).

### 3.4 Nicotinamide has Differential Effects on I_Na_ between HEK293 Cells and RNCMs

NAM (Figure 4A), an alternate NAD^+^ precursor to NR known to inhibit sirtuins at high dose, had no significant effect on I_Na_ in HEK293 cells transiently-transfected with Na_V_1.5 (500 µM: −19 ± 26%, p=0.56; 5 mM: +21 ± 26%, p=0.49; Figure 4B-C, Table S3). No significant changes were observed in steady-state activation and inactivation after NAM administration (Figure S8). Interestingly, NAM stimulated I_Na_ in RNCMs at a lower dose (500 µM: 84 ± 33%, p=0.003) but had no significant effect on I_Na_ at a higher dose (5 mM: −11 ± 24%, p=NS; Figure 4D-E, Table S3). Gating properties were unchanged (Figures 4F, S9, Table S5). Unlike NR, NAM had no effect of I_Na,L_ (Figure 4G). Measurement of NAD^+^ and NADH showed that NAM increased both NAD^+^ and NADH content in a dose-dependent manner with a significant increase in the overall NAD^+^/NADH ratio (500 µM: 7.5 ± 2.0%, p=0.007, 5 mM: 23.8 ± 3.9%, p<0.0001; Figure 4H).

**Figure 4.**
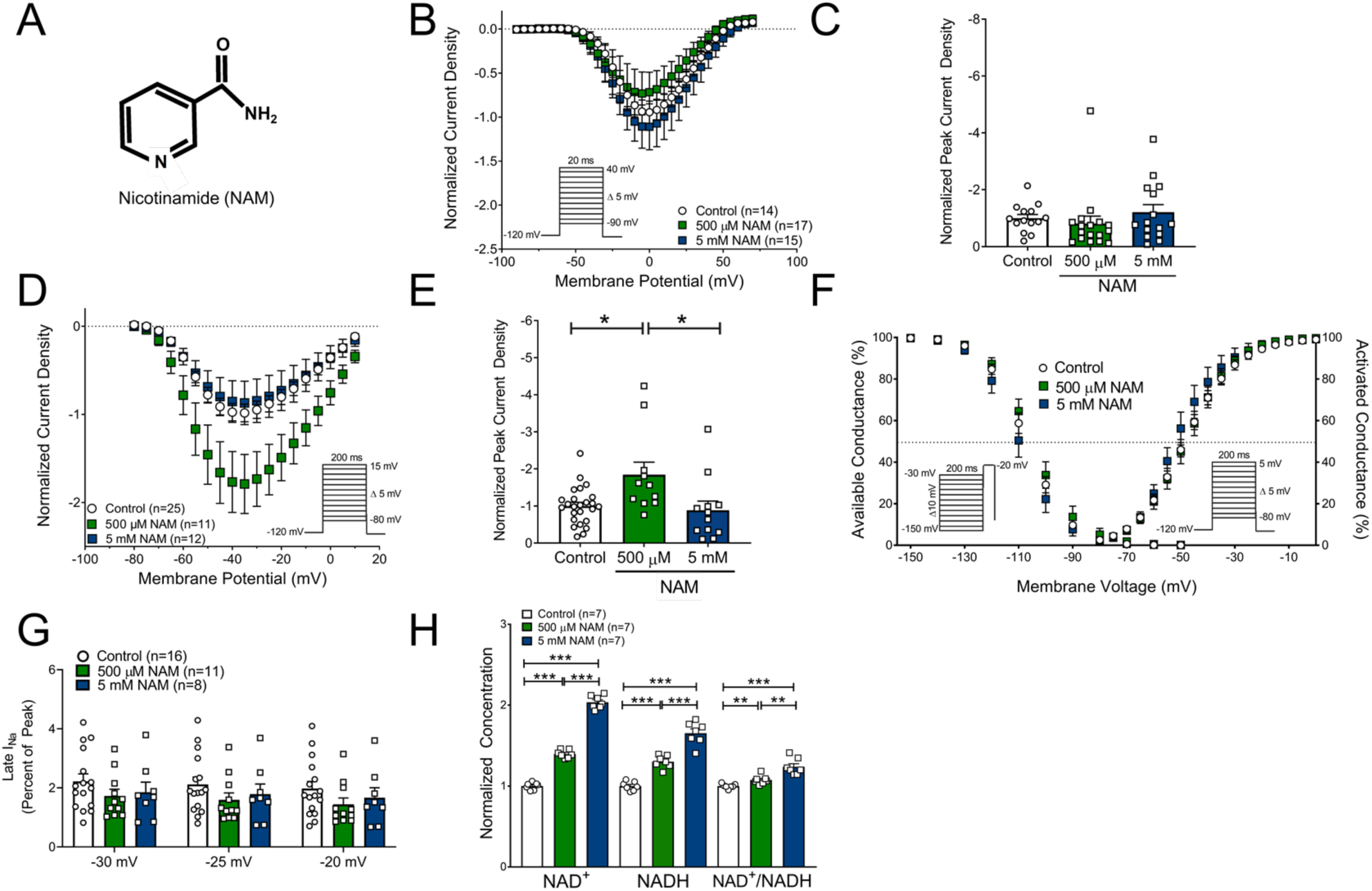
Nicotinamide (NAM), an alternative NAD^+^ precursor, does not modulate Na_V_1.5 in HEK293 cells transiently transfected with Na_V_1.5 but dose-dependently modulates Na_V_1.5 in RNCMs. **(A)** Structure of NAM, **(B)** I-V curve, and **(C)** normalized peak current density of HEK293 cells transiently transfected with Na_v_1.5 and administered NAM (500 µM or 5 mM, 48 hours). **(D, E)** The effect of NAM supplementation (500 µM or 5 mM, 24 hours) on I-V curve and normalized peak current density illustrates dose-dependent effects on Na_V_1.5 activity in RNCMs. 500 µM NAM stimulates I_Na_ whereas 5 mM NAM has no effect on I_Na_. **(F)** Gating properties of steady-state activation and inactivation. (**G)** Late current, I_Na,L_, in RNCMs treated with NAM **(H)** NAD^+^, NADH, and NAD^+^/NADH ratio quantified in RNCMs after 24-hour NAM treatment. Differences between groups were statistically determined utilizing unpaired t-tests (*p<0.05, **p<0.01, ***p<0.001).

### 3.5 1-methyl-Nicotinamide, an inert Nicotinamide Metabolite, has no effect on I_Na_

1-methyl-Nicotinamide (meNAM), an inert byproduct of NAM degradation and not an NAD^+^ precursor, had no significant effect on I_Na_ (500 µM: −7 ± 14%, p=NS; 5 mM: −5 ± 9.9%, p=NS) channel gating properties, or I_Na,L_ in RNCMs (Figure S10A-G). MeNAM tended to increase NAD^+^ and decrease NADH content in a dose-dependent manner, leading to a significant increase in NAD^+^/NADH ratio (Figure S10H).

### 3.6 NAD^+^ Metabolome Following NAD^+^ Metabolite Supplementation in RNCMs

NAD^+^ metabolite supplementation in cardiomyocytes has been relatively unexplored. The effect of supplementation with NR, NAM or meNAM on the RNCM NAD^+^ metabolome was assessed by LC-MS (Figure 5). Both NR and NAM caused dose-dependent increases in NAD^+^ whereas the inert meNAM had no effect on NAD^+^ content (Figure 5A). NR was superior to NAM in increasing NAD^+^ and nicotinamide adenine dinucleotide phosphate (NADP^+^) at equimolar concentrations (Figure 5A,B). However, NAM was superior in increasing nicotinic acid adenine dinucleotide (NAAD, Figure 5C). NR markedly depressed levels of ADP-ribose (ADPR), a substrate for another class of NAD^+^-consuming enzymes, Poly-ADP-Ribose Polymerases (PARPs, Figure 5I). NAM, an inhibitor of PARPs, had no <u>significant</u> effect on ADPR levels.

**Figure 5.**
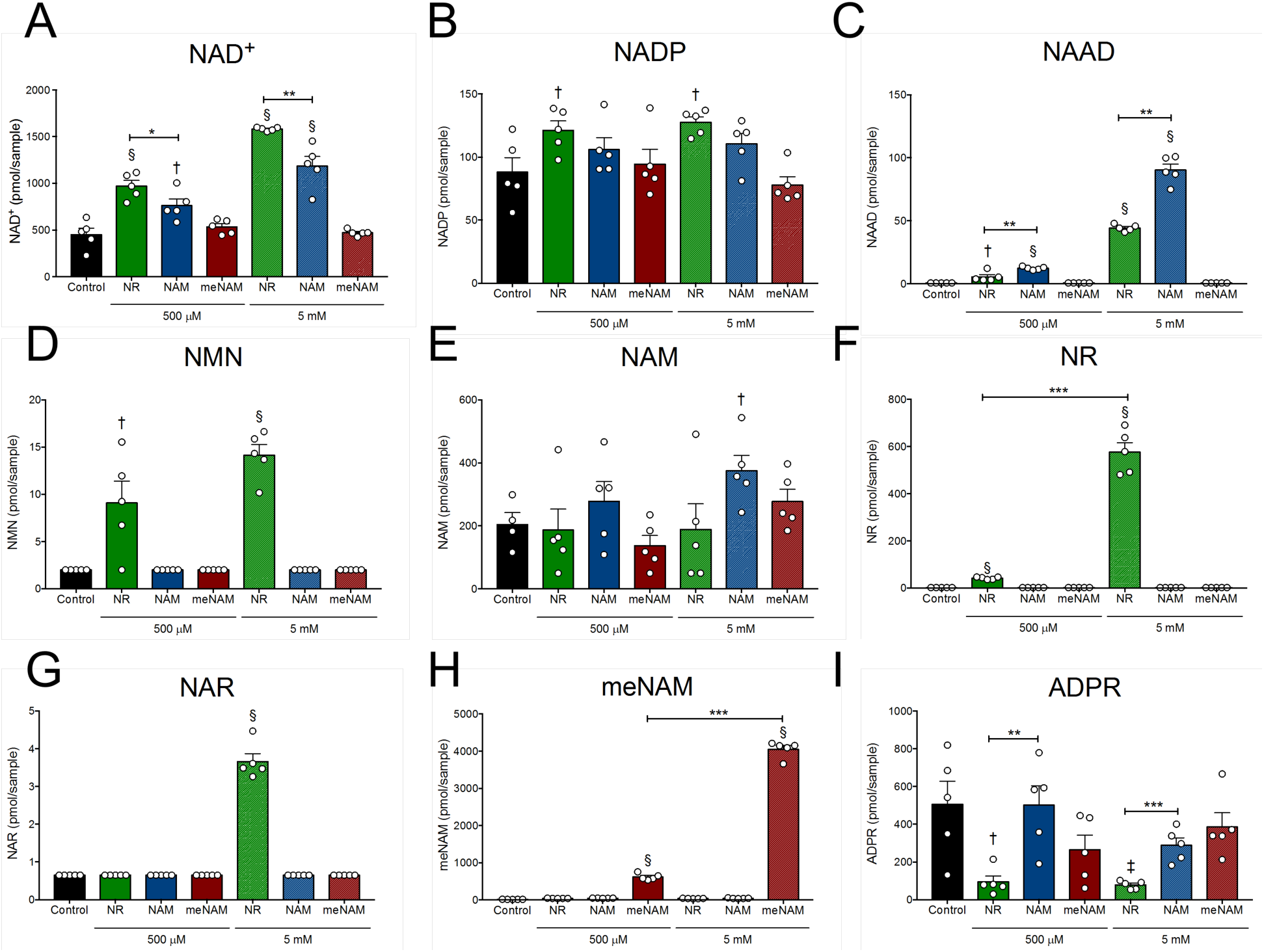
NAD^+^ precursors Nicotinamide Riboside (NR) and Nicotinamide (NAM) increases NAD^+^ Content in Neonatal Rat Cardiomyocytes. **(A-I)** NAD^+^, NADP, NAAD, NMN, NAM, NR, NAR, meNAM, and ADPR were assessed by LC-MS in cellular extracts of neonatal rat cardiomyocytes treated with either H_2_O, NR, NAM, or meNAM (500 µM or 5 mM, 24 hours). Differences between groups were statistically determined utilizing unpaired t-tests (*p<0.05, **p<0.01, ***p<0.001 vs. respective groups and ^†^p<0.05, ^‡^p<0.05, ^§^p<0.001 vs. control).

### 3.7 Disruption of the Na_V_1.5-K1479 Acetylation Site and Na_V_1.5-S1503 Phosphorylation sites Modulates the Effects of NAD^+^ Precursors on I_Na_

SIRT1-dependent deacetylation of Na_V_1.5 in the intracellular Domain III-IV linker at K1479 increased I_Na_ by enhancing membrane trafficking, and disruption of the acetylation by site-directed mutagenesis (Na_V_1.5-K1479A) eliminated the effect of SIRT1 on I_Na_ in HEK293 cells.^15^ To determine the role of acetylation following treatment with NAD^+^ precursors, HEK293 cells transiently transfected with wild-type Na_V_1.5 or Na_V_1.5-K1479A channels were treated with NR or NAM. No differences in the magnitude and gating properties of I_Na_ were observed between the wild-type and K1479A Na_V_1.5 channels under basal conditions (Figure S11). Similar to its effect on wild-type channels (Figure 1G-H), NR increased I_Na_ in HEK293 cells expressing Na_V_1.5-K1479A (5 mM: 75 ± 29%, p=0.02; Figure 6A,B) and significantly shifted steady-state activation and inactivation to more hyperpolarized potentials (Figures 6C, S12, Table S5). However, NR had no effect on I_Na_ in HEK293 cells expressing Na_V_1.5-K1479A/S1503A (5 mM: 16 ± 12%, p=0.38; Figure 6D,E) while maintaining the hyperpolarizing shift in steady-state activation and inactivation (Figures 6F, S12, Table S5). Therefore, disruption of the PKC site abolished the effect of NR on increasing I_Na_.

**Figure 6.**
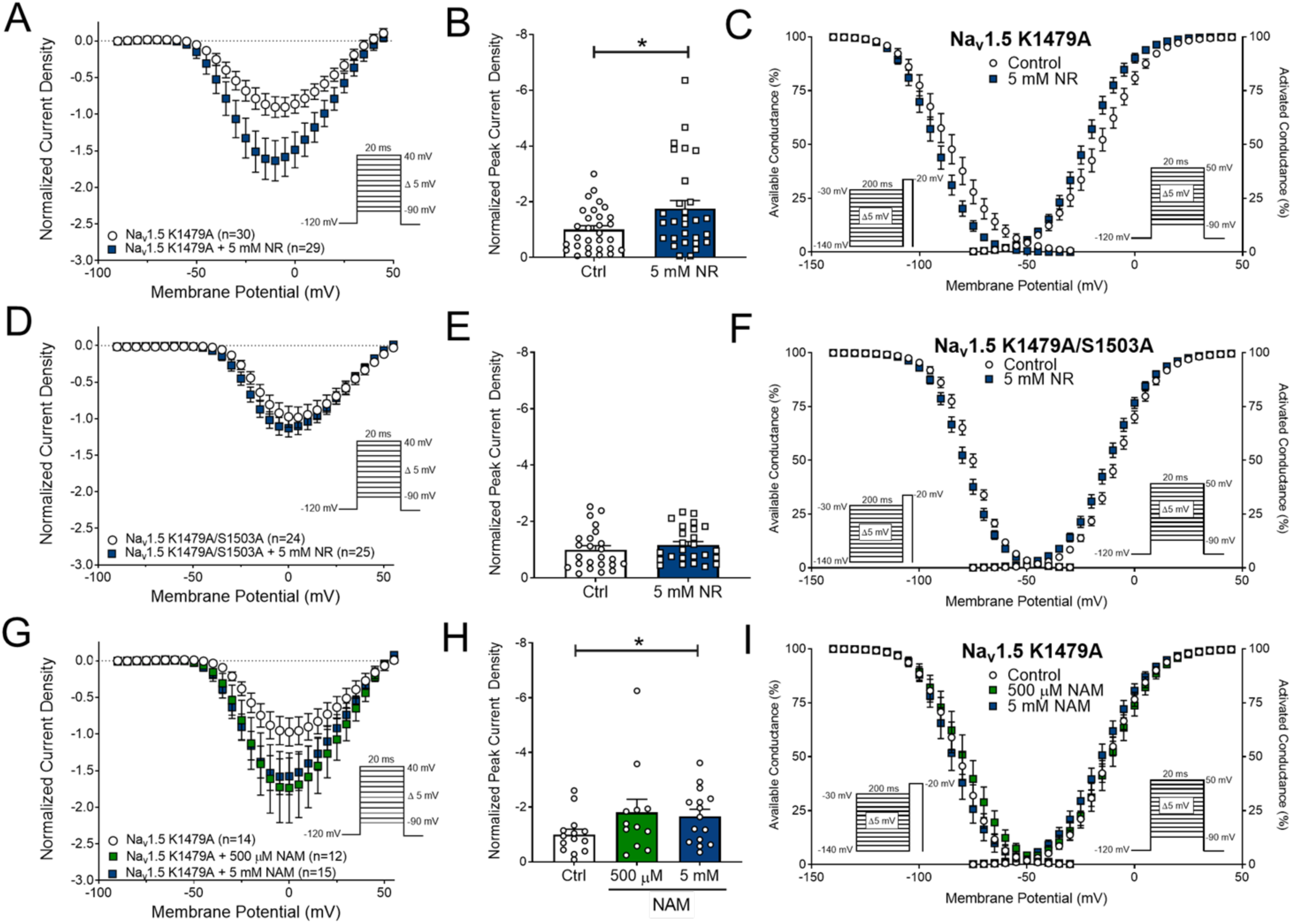
The Role of the Na_V_1.5 Acetylation K1479 and Phosphorylation S1503 sites in mediating the effect of NAD^+^ precursors. The I-V relationship, peak current density, and steady-state activation and inactivation of HEK293 cells transiently transfected with: **(A-C)** the mutant Na_V_1.5 K1479A channel with and without the administration of NR (5 mM, 48 hours), **(D-F)** the mutant Na_V_1.5 K1479A/S1503A channel with and without the administration of NR (5 mM, 48 hours), and **(G-I)** the mutant Na_V_1.5 K1479A channel with and without the administration of NAM (500 µM or 5 mM, 48 hours). Differences between normalized peak current densities were statistically determined utilizing unpaired t-tests (*p<0.05).

Administration of NAM to HEK293 cells expressing Na_V_1.5-K1479A increased I_Na_ (500 µM: 81 ± 47%, p=0.10; 5 mM: 66 ± 25%, p=0.04; Figure 6G-I), unlike our prior finding that NAM had no effect on HEK293 cells expressing wild-type Na_V_1.5 (Figure 4). NAM had no effect on steady-state activation or inactivation (Figure S8). Thus, elimination of the SIRT1-deacetylation site allowed NAM to increase I_Na_.

Together these results suggest that both SIRT1-dependent and SIRT1-independent mechanism are modulating Na_V_1.5 in response to NAD^+^ metabolites that increase NAD^+^.

### 3.8 Dietary Supplementation of Nicotinamide Riboside Decreases QTc in Wild-type Mice

To assess for changes in cardiac electrophysiology with NAD^+^ supplementation *in vivo*, C57BL/6J mice were either placed on a diet with (0.35% or 1.0% by weight) or without (control) supplemental NR. Over the time course of dietary supplementation, no differences in body weight were observed demonstrating that the NR diets are tolerated and did not alter growth (Figure S13). Electrophysiological properties were assessed by high-resolution EKG (Figure 7A-B). No differences in PR and QRS intervals were observed between groups at different timepoints (Figure 7C-D, Table S6). However, mice supplemented with NR demonstrated a decrease in QTc (0.35% NR: −4.9 ± 2.0%, p=0.22, 1.0% NR: −9.5 ± 2.8%, p=0.01) after 10-12 weeks compared to baseline, with no change in mice on a control diet (+1.8 ± 2.3%, p=0.99, Figure 7A,B,E, Table S6). NR supplementation had no effect on cardiac function as assessed by echocardiography (Table S7).

**Figure 7.**
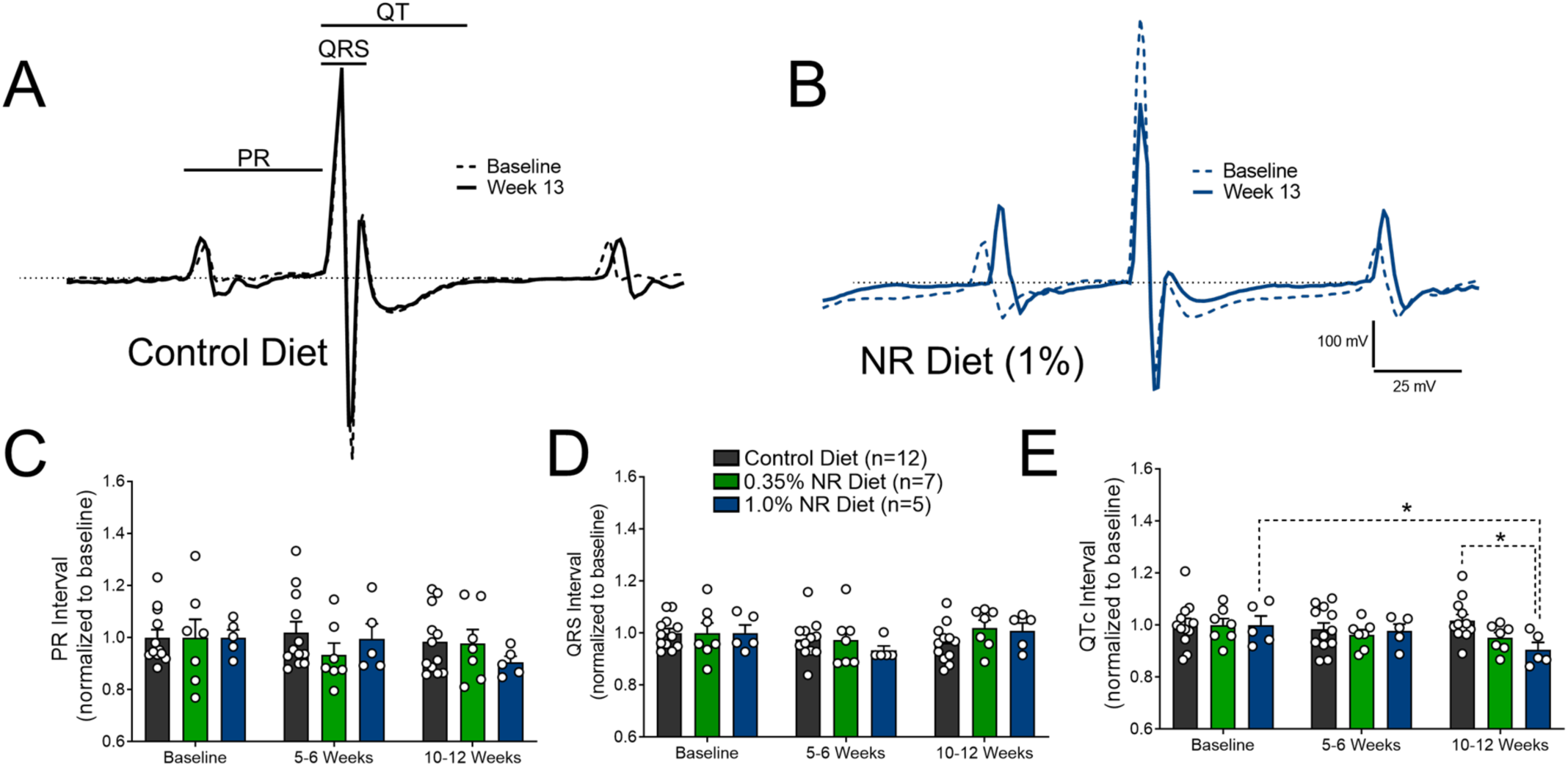
Dietary Supplementation of Nicotinamide Riboside Modulates Cardiac Repolarization. 4-6 month old wild-type C57BL/6 mice were randomly placed on either a control diet or diet supplemented with NR (0.35% or 1%). Cardiac electrophysiological changes were assessed by EKG. **(A, B)** Representative Lead II EKG traces overlaying baseline (dashed line) and 10-12 weeks post-diet (solid line) for mice on either control diet or diet supplemented with 1.0% NR. **(C)** PR interval, **(D)** QRS interval, and **(E)** QTc duration were measured. Significance between groups and within a group at each timepoint was determined by Two-way ANOVA with Bonferroni post-hoc multiple-comparisons test. (*p<0.05).

## 4. DISCUSSION

Our group and others have explored the molecular basis for the regulation of the cardiac sodium channel Na_V_1.5 by NAD^+^, NADH, and the NAD^+^/NADH ratio. Previous studies demonstrated that exogenous application of 100 μM NAD^+^ on Langendorff-perfused hearts from *Scn5a* haplo-insufficient mice was protective against programmed electrical stimulation induced polymorphic ventricular tachycardia.^12^ In addition, exogenous *in vitro* application and *in vivo* intraperitoneal injection of NAD^+^ rescued the decrease in peak I_Na_ in myocytes from mice with a non-ischemic cardiomyopathy (with hypertension from unilateral nephrectomy, deoxycorticosterone acetate pellet implantation, and salt water substitution).^25^ Furthermore, application of NAD^+^ on failing human heart wedge preparations improved conduction velocity.^25^

In this study, we demonstrate that NR, a commercially bioavailable NAD^+^ precursor,^26^ can increase peak I_Na_ in heterologous expression systems and RNCMs without influencing the NAD^+^/NADH ratio. We also show, for the first time, that NAD^+^ precursor supplementation reduces I_Na,L_ in RNCMs and shortens QTc in mice, collectively suggesting that both an increase in peak I_Na_ and a decrease in I_Na,L_ are novel potential antiarrhythmic mechanisms following NAD^+^ supplementation. Together, these findings provide substantial evidence for a direct role of NAD^+^ in regulating the electrical activity of the heart through modulation of Na_V_1.5.

Translation and trafficking of Na_V_1.5 to the cellular membrane surface is tightly regulated by a series of processes involving microRNA-based mRNA degradation, translation, assembly, transport through the endoplasmic reticulum and Golgi complex, anchoring at the cellular surface, recycling of the channel through internalization, and degradation.^27, 28^ A variety of mechanisms coordinate the trafficking of Na_V_1.5 and involve a collection of accessory interacting proteins and post-translational modifications. At the surface, Na_V_1.5 function can be dictated by open probability, conductance, and gating properties. We previously reported that acetylation of Na_V_1.5 at a highly conserved lysine (K1479) in the Domain III-IV intracellular linker, a region of the channel historically known for its role in channel inactivation, regulates channel surface expression but not gating properties.^15^ However, the mechanisms regulating Na_V_1.5 amplitude and properties in response to NAD^+^ precursor supplementation remain unknown.

Our data here demonstrates that the application of 500 µM to 5 mM of the NAD^+^ precursor NR increased peak I_Na_ by approximately 40-60% in HEK293 cells stably-expressing Na_V_1.5, HEK293 cells transiently-transfected with Na_V_1.5, and RNCMs with no significant change in Na_V_1.5 expression, channel steady-state gating properties, or channel surface and membrane expression. We also show that NR decreases K1479 Na_V_1.5 acetylation only at the higher 5 mM dose. Together, these findings suggest suggestive that the increase in I_Na_ observed at both doses (500 µM and 5 mM) is independent of acetylation in heterologous cell systems are cardiac myocytes.

NAD^+^ is utilized as a substrate for sirtuins (SIRTs) and activity is dependent on NAD^+^ levels. NR drives sirtuin activity by increase NAD^+^ content. SIRT1 (Sirtuin 1), the primary and most evolutionary conserved sirtuin, deacetylates Na_V_1.5 at K1479 in heterologous cell systems, RNCMs, and mice.^15^. The enzymatic reaction of sirtuin deacetylation results in the breakdown of NAD^+^ to NAM and ADP-ribose. NAM also holds the potential to regenerate NAD^+^ but at high doses can be a competitive feedback inhibitor on sirtuins and other NAD^+^-consuming enzymes, and we have previously shown that 5 mM NAM increases K1479 acetylation.^15^ In this study, the application of NAM on RNCMs demonstrated a dose-dependent effect on I_Na_, with low doses stimulating I_Na_ but higher doses having no effect on I_Na_ despite increasing NAD^+^ content and NAD^+^/NADH ratio. This illustrates the effects of NAM in boosting NAD^+^ synthesis but inhibitory of sirtuins. NAM at either dose (500 μM or 5 mM) had no effect of I_Na_ in HEK293 cells transiently transfected with wild-type Na_V_1.5. Furthermore, we confirmed the importance of acetylation in modulating the response of I_Na_ to NAM using the Na_V_1.5 K1479A mutant. The absence of the Na_V_1.5-K1479 acetylation site did not significantly blunt the effect of NR on I_Na_, suggesting that sirtuin-mediated deacetylation is not primarily responsible for the increase on I_Na_ following NR administration. However, disruption of the acetylation site enabled NAM to increase I_Na_ in HEK293 cells transiently-transfected with Na_V_1.5-K1479A. This illustrates that NAD^+^ metabolites can modulate Na_V_1.5 through both sirtuin-dependent and sirtuin-independent mechanisms.

In addition to acetylation, both protein kinase C (PKC) and protein kinase A (PKA) phosphorylation of Na_V_1.5 has been identified as being sensitive to NAD^+^, NADH, and metabolic stress. Increased levels of NADH activate PKC, resulting in the production of mitochondrial reactive oxygen species (ROS) and the direct phosphorylation of Na_V_1.5 at S1503 within the Domain III-IV linker region.^13^ These actions have been shown to decrease single channel conductance as well as decrease channel surface expression.^13, 14^ By increasing NAD^+^ content, the production of reactive oxidant species is reduced and therefore, the application of NR may modulate the PKC-mediated phosphorylation and oxidant stress production in the regulation of Na_V_1.5.^25^ Our data demonstrates that the disruption of the PKC site at S1503 abolished the effect of NR, suggesting that the effect of NR on increasing I_Na_ is dependent on the PKC phosphorylation site, which is known to decrease single channel conductance. As opposed to PKC, the stimulation of PKA phosphorylation, shown to be mediated by NAD^+^, blunts the effects of NADH on decreasing I_Na_.^12, 29, 30^ PKA stimulation has been shown to have no effect on I_Na,L_ in wild-type Na_V_1.5 channels.^31^ Together, the interplay between acetylation, phosphorylation, and mitigation of oxidant stress may provide an explanation of NAD^+^-mediated modulation of peak I_Na_ through the cardiac sodium channel Na_V_1.5 (Figure 8).

**Figure 8.**
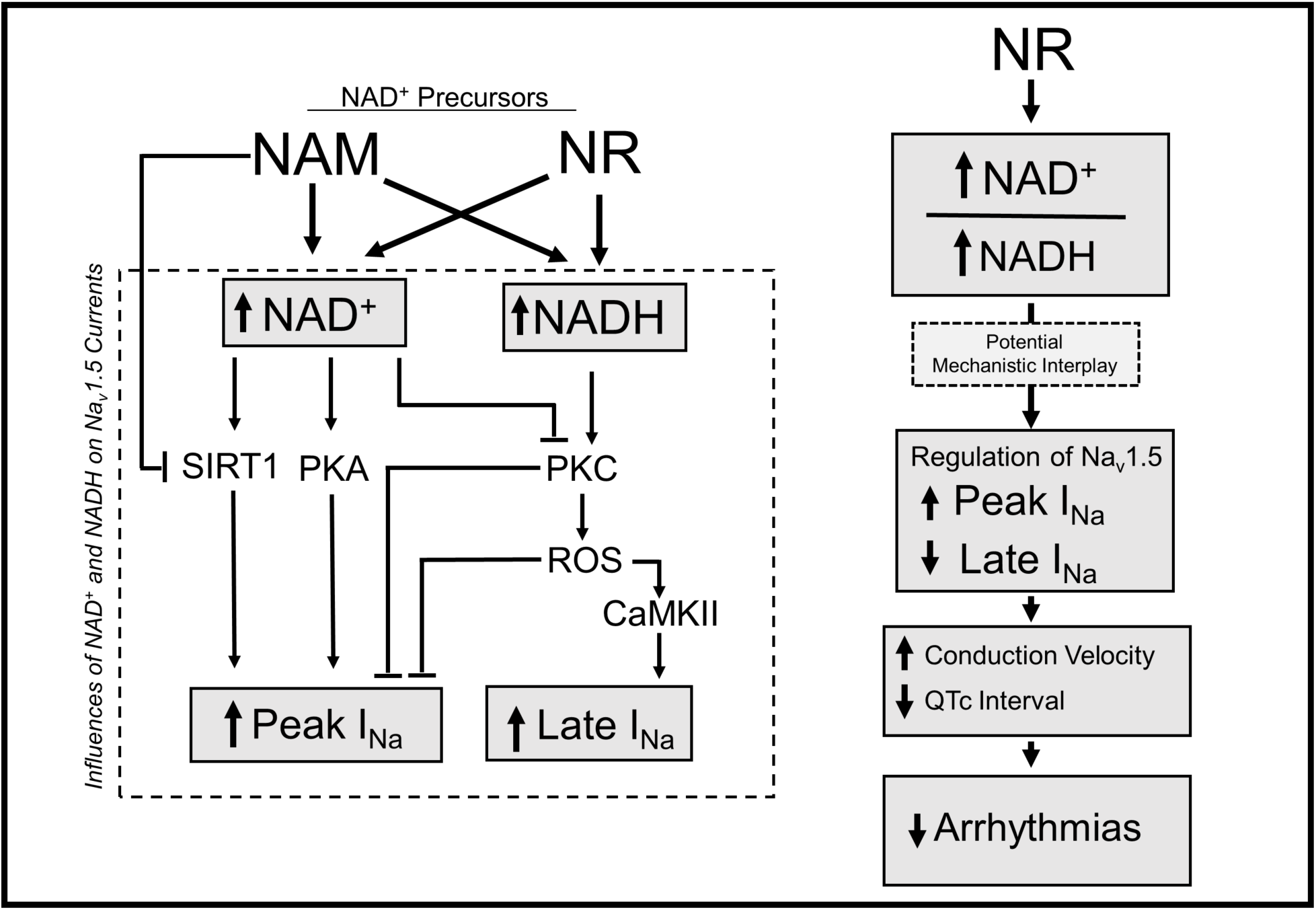
Model of NAD^+^ Precursor Modulation of Na_V_1.5.

NAD^+^ supplementation may also drive other unidentified post-translational modifications of Na_V_1.5 including ADP-ribosylation. In addition to sirtuins, NAD^+^ is consumed as a substrate by poly(ADP)-ribose polymerases (PARPs) to catalyze the addition of ADP-ribose moieties onto target proteins.^32^ Our data illustrates that NAD^+^ supplementation with NR may be stimulating ADP-ribosylation, as ADP-ribose levels are markedly depressed after NR treatment in RNCMs (Figure 5). Of note, similar decreases of ADP-ribose are not observed after NAM treatment as NAM inhibits PARP activity.^33^ Investigation of ADP-ribosylation may provide further insight into the NAD^+^-dependent regulation of Na_V_1.5.

Previous reports have indicated that CD38 mediates the effect of extracellular NAD^+^ in rescuing I_Na_ in response to increased intracellular NADH.^25^ CD38 has been shown catalyze intracellular NAD^+^ to cyclic ADP-ribose in addition to metabolizing extracellular NAD^+^ precursors.^34^ Although HEK293 cells do not express CD38, HEK293 cells and fetal bovine serum have been shown to metabolize NAD^+^ precursors into other intermediates.^35^ In our *in vitro* studies, media and treatment was replenished every 24 hours to prevent precursor degradation over the time course of the experiment. Collectively, CD38 activity and other NAD^+^ intermediates may play additional roles in regulating the cardiac sodium channels and cardiac electrophysiology that are not yet known.

An increase in the sustained or late cardiac Na^+^ current, I_Na,L_, seen in congenital long QT syndrome type 3 (LQT3) and in acquired disorders including cardiomyopathies, can induce a pro-arrhythmic state through prolongation of the action potential leading to early afterdepolarizations while also contributing to myocyte sodium and calcium loading.^36–38^ Blockers of I_Na,L_ including flecainide in LQT3 and ranolazine have been studied as potential antiarrhythmic agents.^39–41^ Our data demonstrates a reduction in I_Na,L_ with NR treatment in RNCMs *in vitro* and QTc shortening after NR supplementation in mice *in vivo*. A potential mechanism of NR reducing I_Na,L_ may be mitigating oxidative stress. Oxidative stress and reactive oxygen species (ROS) decrease I_Na_ and increase I_Na,L_, contributing to electrical and contractile dysfunction in heart failure.^42, 43^ The activation of Ca/Calmodulin-Kinase II (CaMKII) and the CaMKII-dependent phosphorylation of Na_V_1.5 is required for oxidative stress to increase I_Na,L_.^44^ NR supplementation and increased NAD^+^ metabolites may decrease oxidative stress and ROS, leading to lower CaMKII activity and decreased I_Na,L_. NR may have additional effects on other ion channels and their subunits including voltage-gated K^+^ and Ca2^+^ channels that mediate repolarizing outward potassium currents and QTc.^45^ We did not see changes in K^+^ or Ca^2+^ currents in RNCMs following NR supplementation that would shorten action potential duration and QTc. It is important to note the differences between mouse and human cardiac electrophysiology as mouse action potentials are much shorter and regulated by different repolarizing K^+^ currents.^46^

Modulating Na_V_1.5 provides the therapeutic potential for preventing arrhythmias in a variety of inherited and acquired diseases associated with Na_V_1.5 dysfunction including Brugada Syndrome, sick sinus syndrome, progressive conduction disease, LQT3 and ischemic/nonischemic cardiomyopathies. Our data demonstrates the ability of NR to increase peak I_Na_ while reducing I_Na,L_ in heterologous systems and RNCMs. The potential relevance of these findings was reinforced by QTc shortening in mice after dietary supplementation of NR. Collectively, these novel findings provide preclinical evidence and the foundation for further testing of select NAD^+^ precursors for the prevention of arrhythmias in conditions associated with Na_V_1.5 dysfunction that decreases I_Na_ and/or increases I_Na,L_.

## 5. CONCLUSIONS

Boosting NAD^+^ content using the NAD^+^ precursor NR but not NAM increased peak I_Na_ in heterologous expression systems and RNCMs through both acetylation-independent and acetylation-dependent mechanisms. In addition, NR supplementation decreased I_Na,L_ in RNCMs and shortened QTc in mice. These findings reinforce the potential role of the NAD^+^ supplementation in cardiac electrophysiology and warrant investigation of NR supplementation as an antiarrhythmic strategy in inherited and acquired disorders associated with abnormal Na^+^ currents.

## 6. ACKNOWLEDGEMENTS

We thank Dr. Samuel Dudley (University of Minnesota) for graciously providing the Na_V_1.5-stably expressing HEK293 cell line and Drs. Christopher Ahern and Daniel Infield for helpful discussions and critical reading of the manuscript. Additionally, we would like to thank Julie S. Jacobs for her technical advice throughout the project.

## 8. Abbreviations

Na_v_1.5: cardiac sodium channel
I_Na_: Na^+^ current
I_Na,L_: late Na^+^ current
SCN5A: cardiac sodium channel gene
PKC: Protein Kinase C
PKA: Protein Kinase A
SIRT1: Sirtuin 1
RNCM: Neonatal Rat Cardiomyocyte
meNAM: N1-*methylnicotinamide*
NAM: Nicotinamide
NR: Nicotinamide Riboside
LQT3: Long QT Syndrome Type 3
CD38: cluster of differentiation 38
PARP: Poly-ADP-Ribose Polymerases
NADP^+^: nicotinamide adenine dinucleotide phosphate
NAD^+^: β-nicotinamide adenine dinucleotide
NADH: reduced form of nicotinamide adenine dinucleotide

## SUPPLEMENTARY DATA

**Figure S1.**
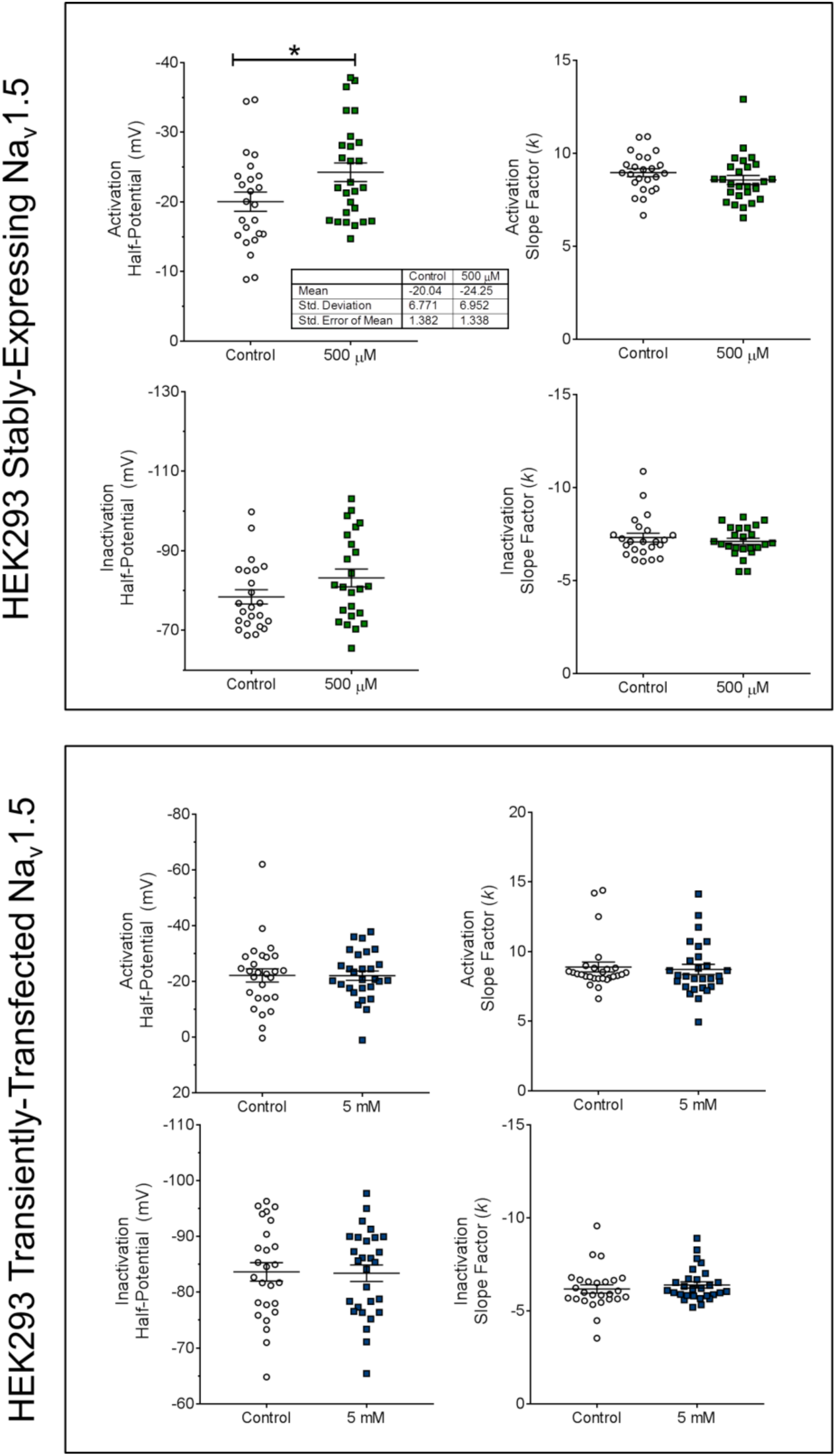
The Effect of Nicotinamide Riboside (NR) on Na_V_1.5 Gating Properties in HEK293 Cells. Na_v_1.5 Gating properties (including half-maximal potentials and slope factors) for steady-state activation and inactivation for **(Top)** HEK293 cells stably-expressing Na_V_1.5 and treated with 500 µM for 48 hours and **(Bottom)** HEK293 cells transiently transfected with Na_V_1.5 and treated with 5 mM NR for 48 hours. Differences between groups were statistically determined utilizing unpaired t-tests (*p<0.05).

**Figure S2.**
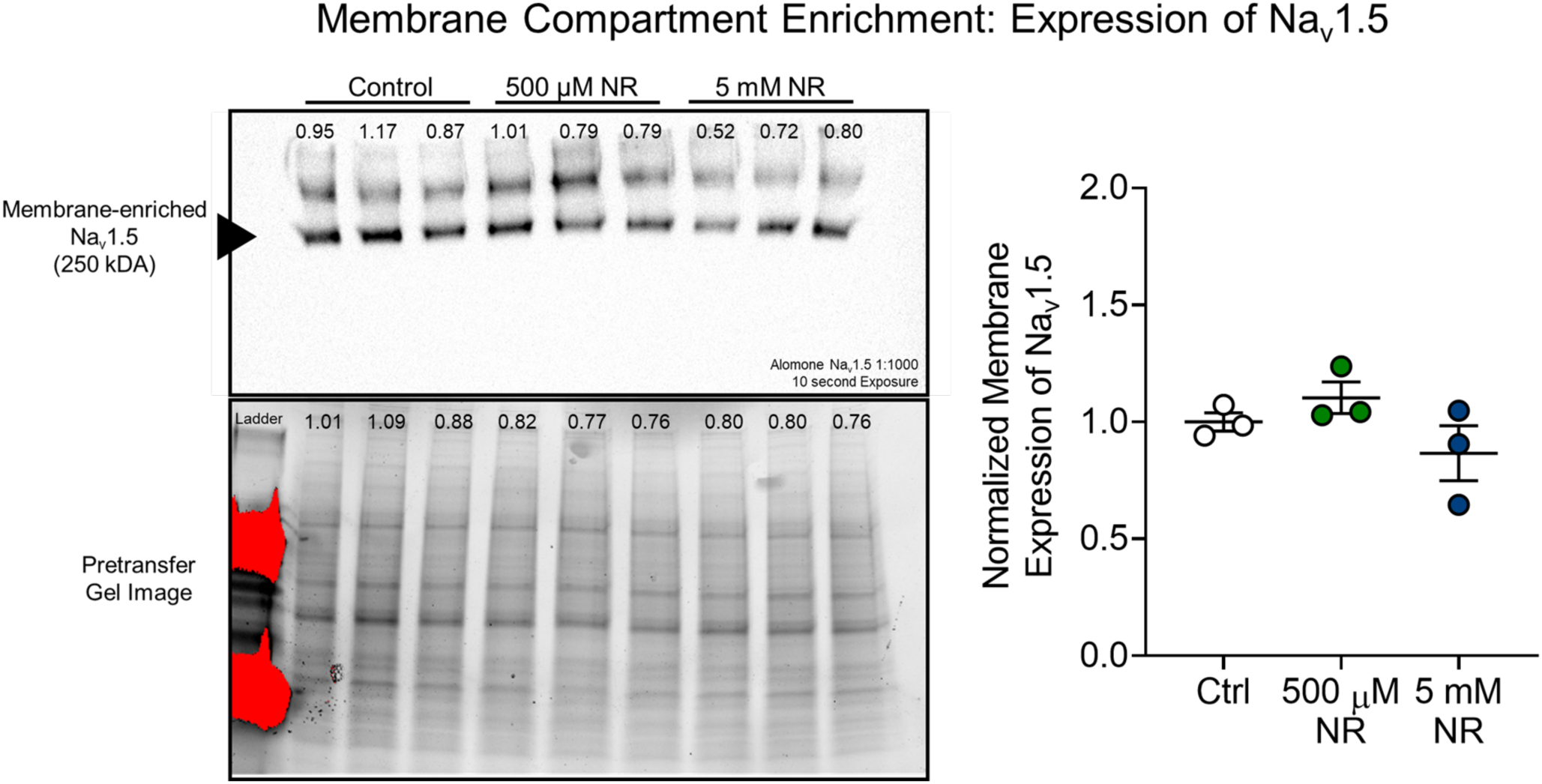
The Effect of Nicotinamide Riboside (NR) on Membrane Expression of Na_v_1.5 in HEK293 Cells stably-expressing Na_v_1.5. HEK293 cells stably expressing Na_v_1.5 were treated with NR (500 µM or 5 mM, 48 hours) and the membrane-enriched protein fraction was harvested with the Compartmentalization Protein Extraction Kit (Millipore, Burlington, MA). Membrane-enriched Na_v_1.5 was blotted, quantified, and normalized to the stain-free gel image obtained prior to transfer to a polyvinylidene fluoride (PVDF) membrane.

**Figure S3.**
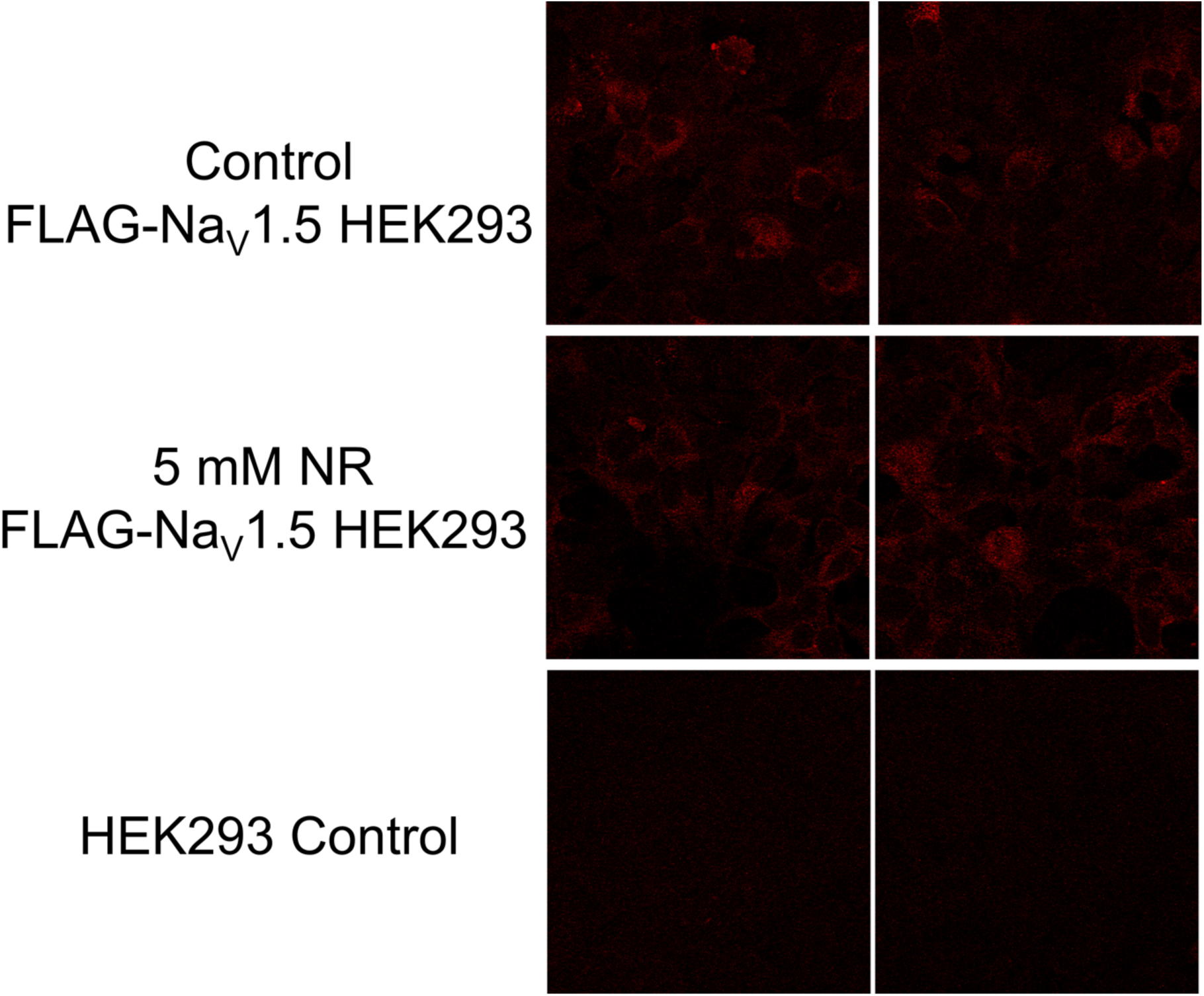
The Effect of Nicotinamide Riboside (NR) on Surface Expression of FLAG-Na_V_1.5. HEK293 cells stably-expressing Na_V_1.5 with a FLAG tag in the extracellular region of D1 was treated with 5 mM NR for 48 hours (replenished at 24 hours) and assessed for surface expression by confocal microscopy. HEK293 cells without Na_V_1.5 were used as a control.

**Figure S4.**
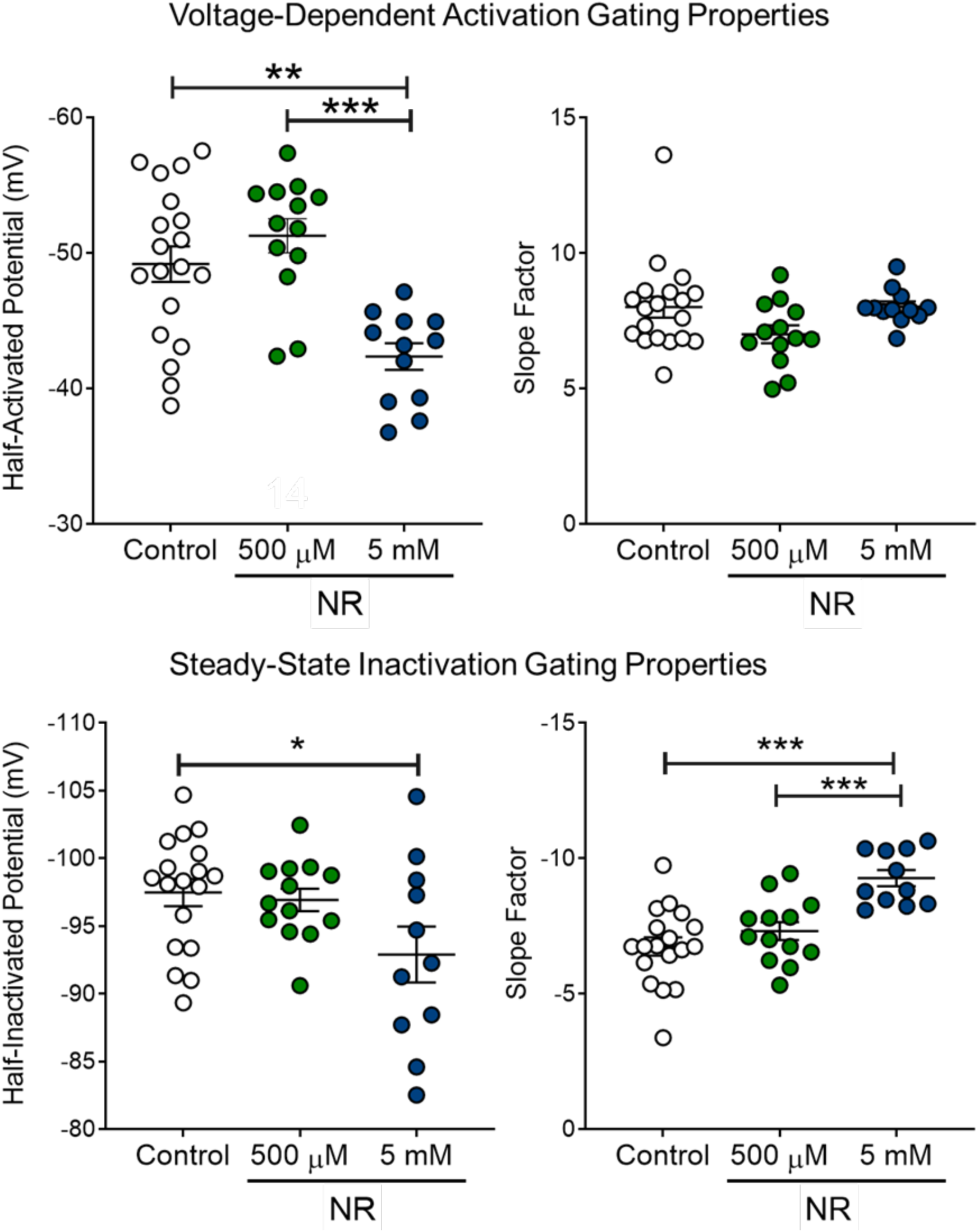
The Effect of Nicotinamide Riboside (NR) on Normalized I_Na_ Gating Properties in RNCMs. (Top) Steady-state activation and **(Bottom)** inactivation properties of neonatal rat cardiomyocytes (RNCMs) treated with H_2_O (vehicle) or NR at either 500 μM or 5 mM for 24 hours. Myocyte properties were normalized to vehicle litter-controlled myocytes to account for litter-to-litter variation in harvesting myocytes. Differences between groups were statistically determined utilizing unpaired t-tests (*p<0.05, **p<0.01, ***p<0.001).

**Figure S5.**
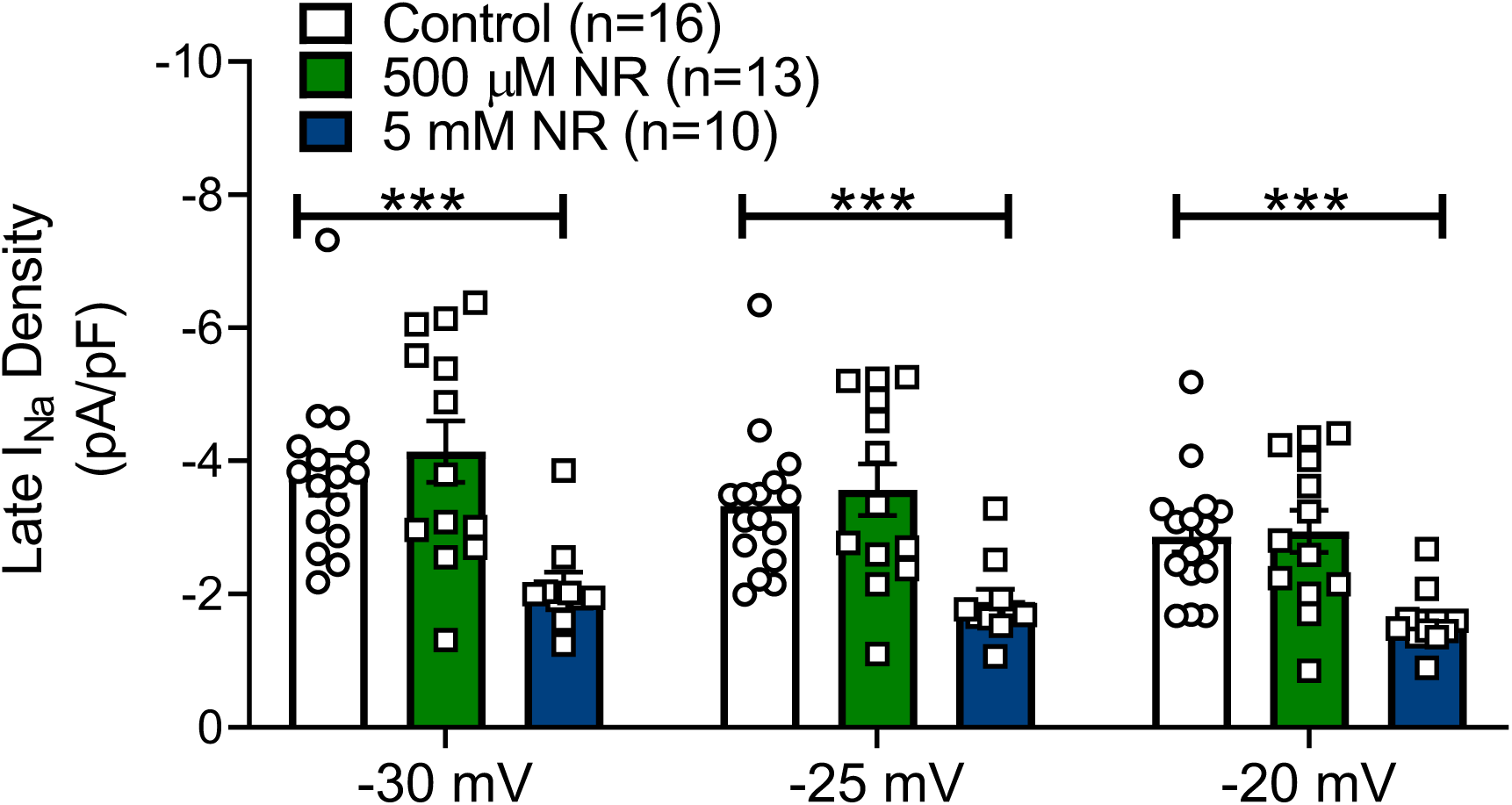
The Effect of Nicotinamide Riboside (NR) on Late Sodium Current Density in RNCMs at. Differences between groups were statistically determined utilizing unpaired t-tests (***p<0.001).

**Figure S6.**
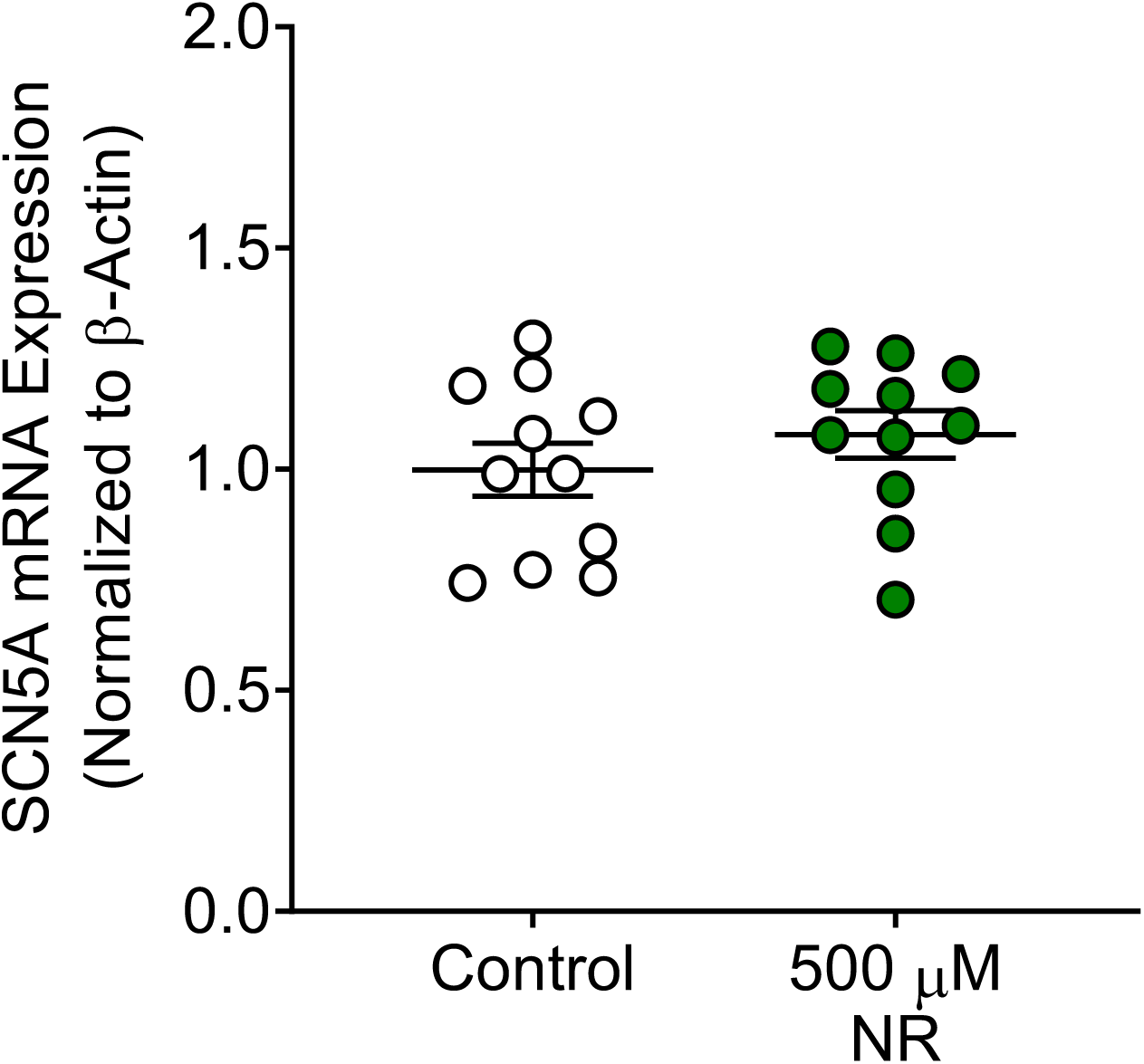
The Effect of Nicotinamide Riboside (NR) on *Scn5a* transcript expression in Neonatal Rat Cardiomyocytes. *Scn5a* transcript expression (normalized to beta-actin expression) was unchanged with administration of 500 μM NR for 24 hours (Ctrl: 1.00 ± 0.05 vs. NR: 1.079 ± 0.05, p=0.33 unpaired t-test).

**Figure S7.**
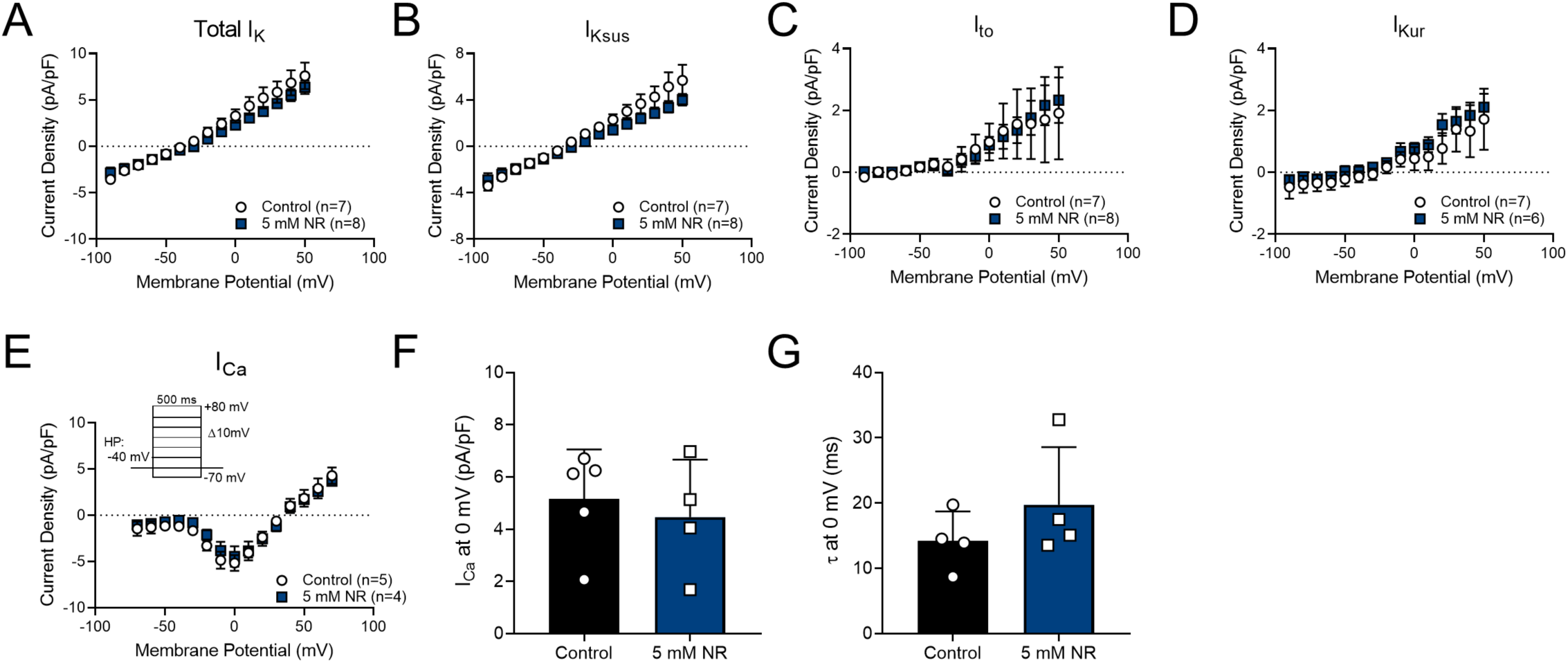
NR has no effect on I_K_ and I_Ca_ Currents. **(A-B)** Whole-cell current-voltage relationships measured at 10 ms into depolarization (I_Ktotal_) and just prior to repolarization to measure the sustained I_K_ (I_Ksus_) recorded from rat cardiomyocytes in control or NR-treated (5 mM, 24 hours) myocytes. **(C-D)** Current-voltage relationship of transient outward K^+^ current (I_to_) and ultra-rapid K^+^ current (I_kur_). No statistically difference between control and NR-treated cells was observed at various membrane potentials. **(E)** Current-voltage relationships for calcium currents recorded from neonatal rat cardiomyocytes. **(F)** The peak current density and **(G)** inactivation time constant (τ) estimated from exponential fits to currents measured at 0 mV recorded from cells control or NR-treated (5 mM NR, 24 hours). Values are mean ± SE.

**Figure S8.**
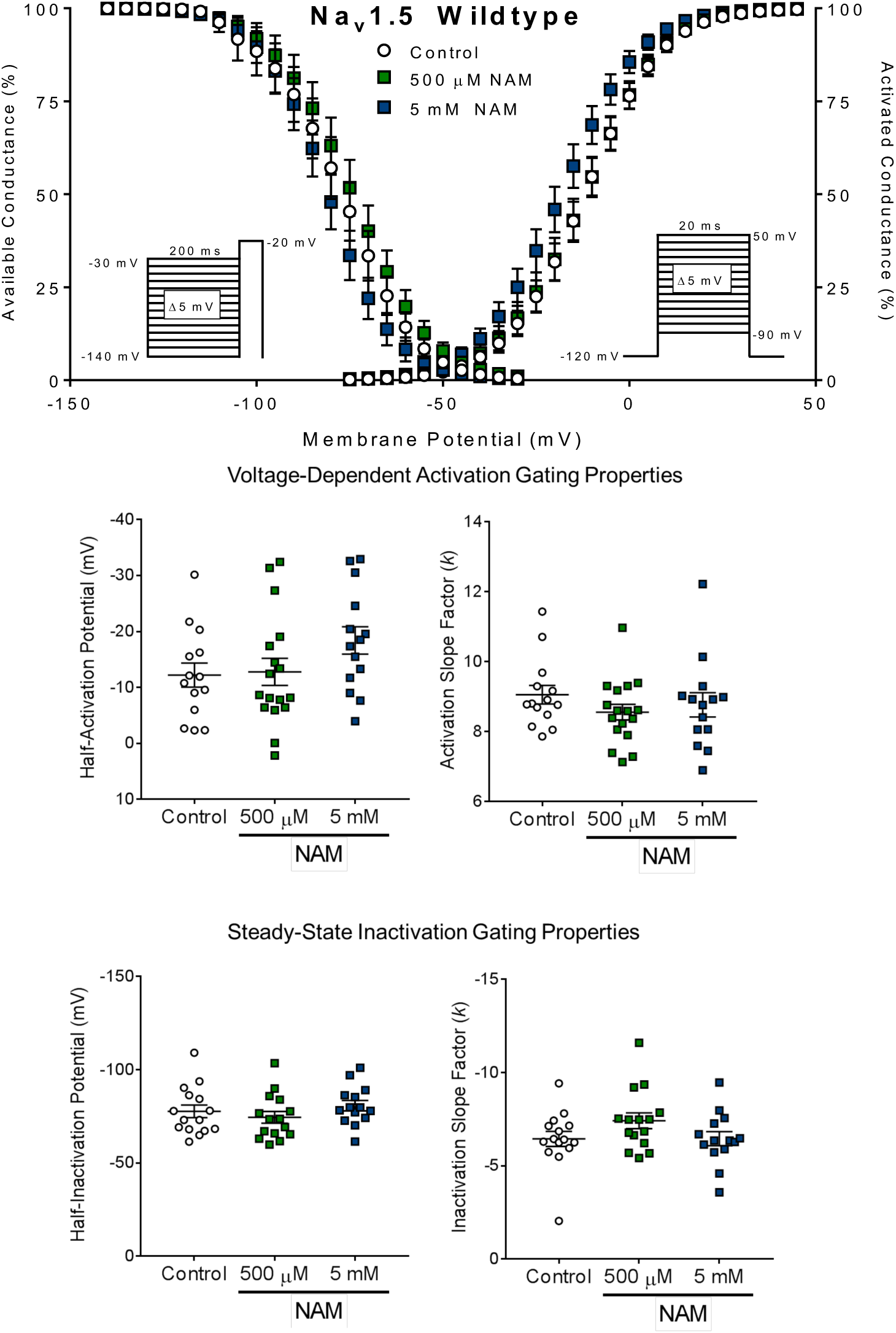
The Effect of Nicotinamide (NAM) on Na_V_1.5 Gating Properties in HEK293 Cells. **(Top)** Gating properties of steady-state activation and inactivation demonstrate are unchanged with NAM treatment. **(Bottom)** Na_V_1.5 gating properties (including half-maximal potentials and slope factors) for steady-state activation and inactivation of HEK293 cells transiently transfected with Na_v_1.5 and treated with H_2_O (vehicle), 500 µM NAM, or 5mM NAM for 48 hours.

**Figure S9.**
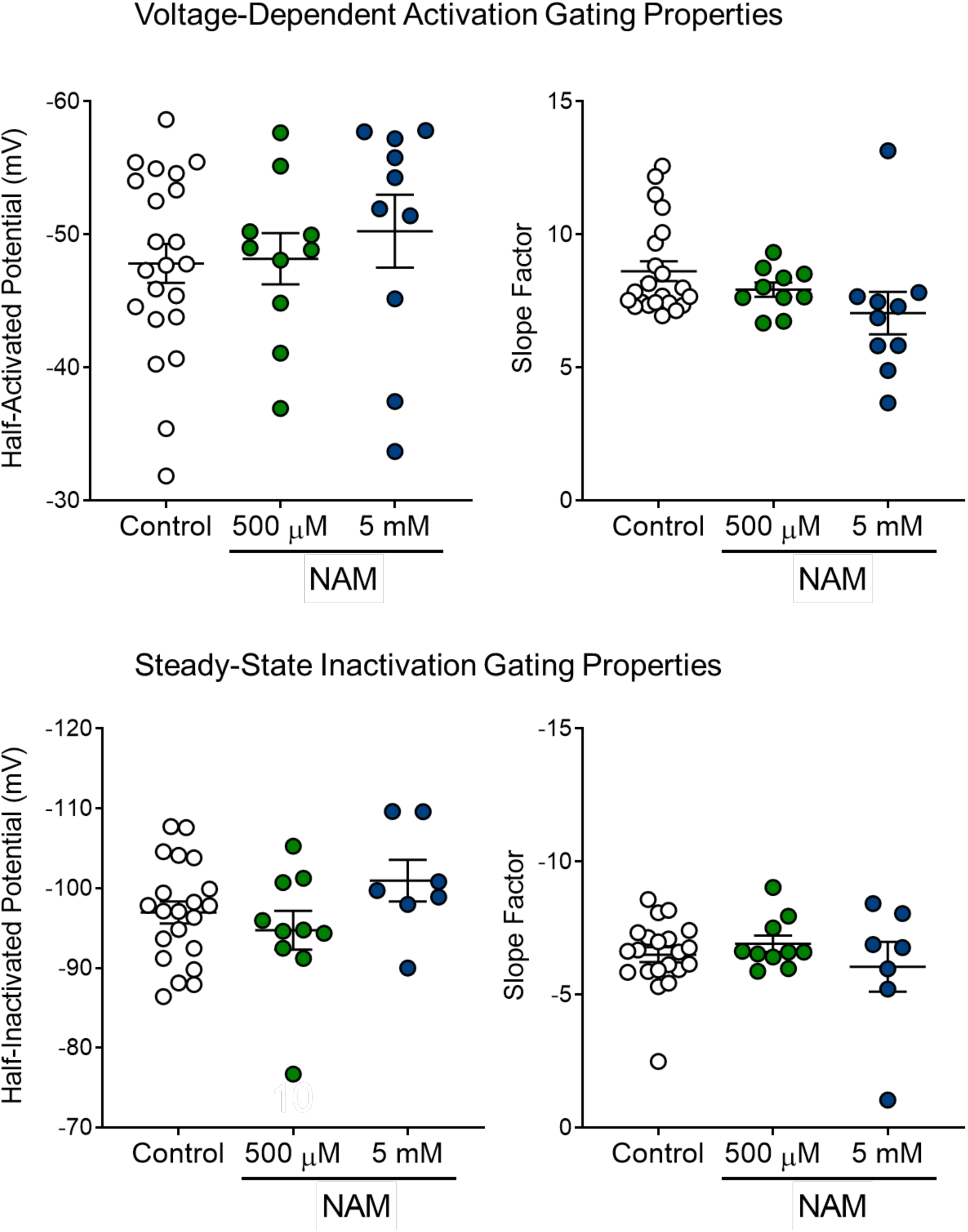
The Effect of Nicotinamide (NAM) on I_Na_ Gating Properties in RNCMs. Steady-state activation and inactivation properties of neonatal rat cardiomyocytes (NRCMs) treated with H_2_O (vehicle) or NAM at either 500 μM or 5 mM for 24 hours. Myocyte properties were normalized to vehicle litter-controlled myocytes to account for litter-to-litter variation in harvesting myocytes.

**Figure S10.**
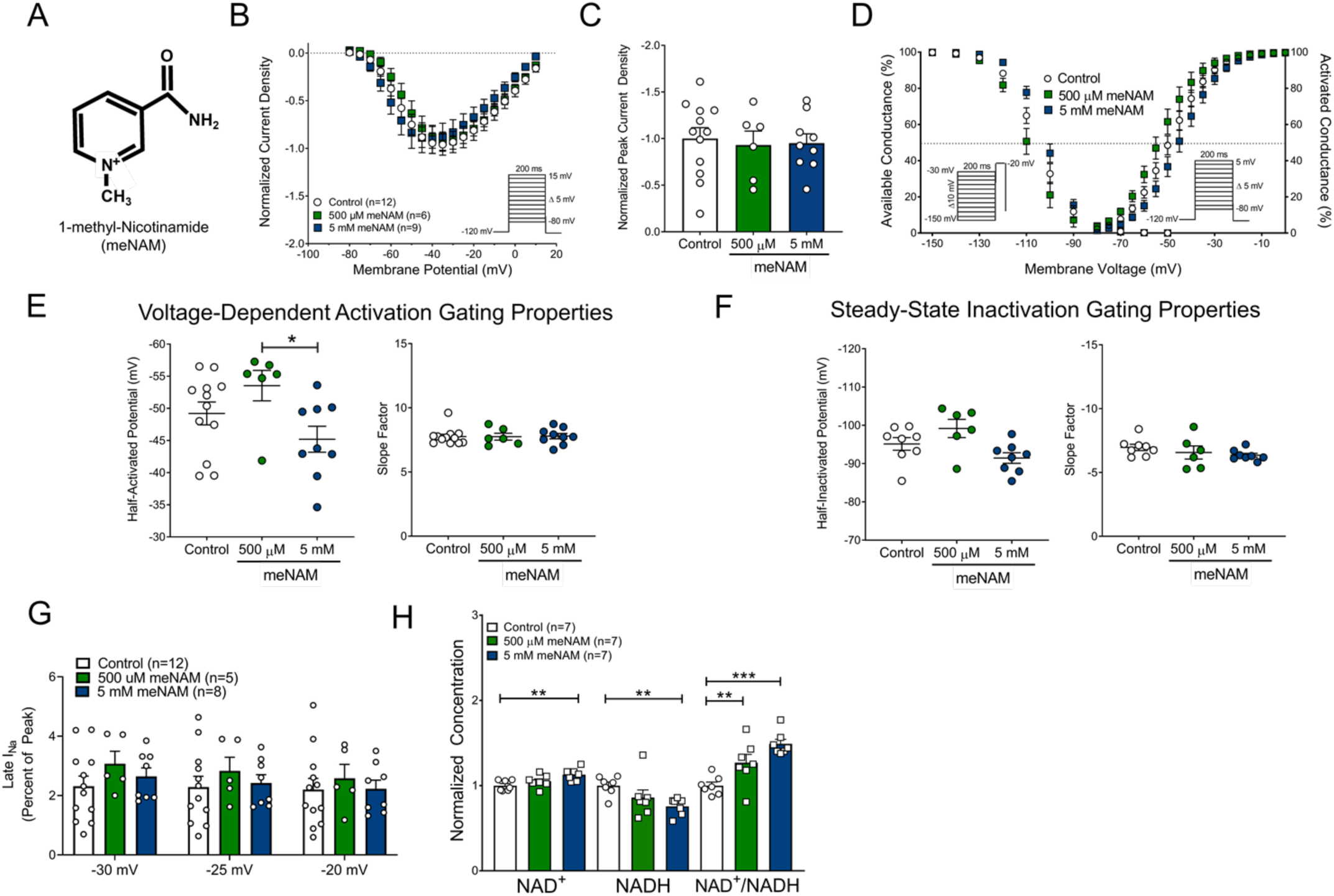
Inert NAD^+^ Metabolite 1-Methyl-Nicotinamide has no effect on I_Na_ in Neonatal Rat Cardiomyocytes. **(A)** The structure of 1-methyl-Nicotinamide (meNAM), an inert byproduct of Nicotinamide which cannot be utilized for NAD^+^ synthesis. **(B,C,D)** meNAM has no to minimal effects on I_Na_ indicated by I-V curve, normalized peak current density, and steady-state activation and inactivation curves. **(E-F)** Gating properties are unchanged after treatment with meNAM. **(G)** Late I_Na_ was unaffected with meNAM supplementation. (**H)** NAD^+^, NADH, and NAD^+^/NADH ratio quantified in rat myocytes after 24-hour meNAM treatment.

**Figure S11.**
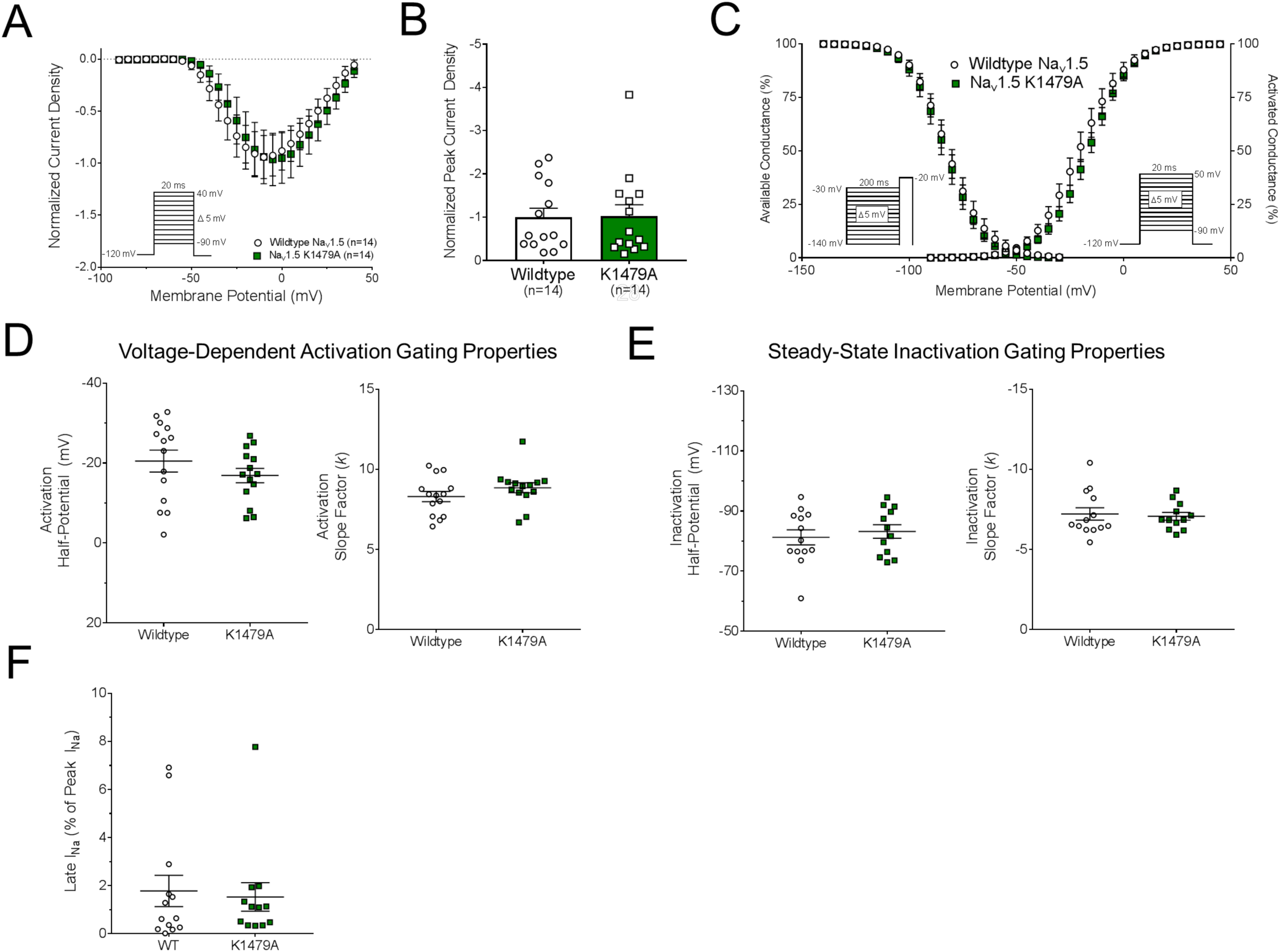
Electrophysiological Comparison of the Wild-type and Mutant K1479A Na_V_1.5 Channel. **(A-C)** The I-V relationship, peak current density, and steady-state activation and inactivation of HEK293 cells transiently transfected with either the wild-type or mutant Na_V_1.5 K1479A channel. **(D-E)** Na_V_1.5 gating properties (including half-maximal potentials and slope factors) for steady-state activation and inactivation of HEK293 cells transiently transfected with wild-type and K1479A Na_V_1.5. **(F)** Comparison of late I_Na_ between wild-type and K1479A Na_V_1.5 as measured by the 10 ms average current at 200 ms post-deploarization at −20 mV.

**Figure S12.**
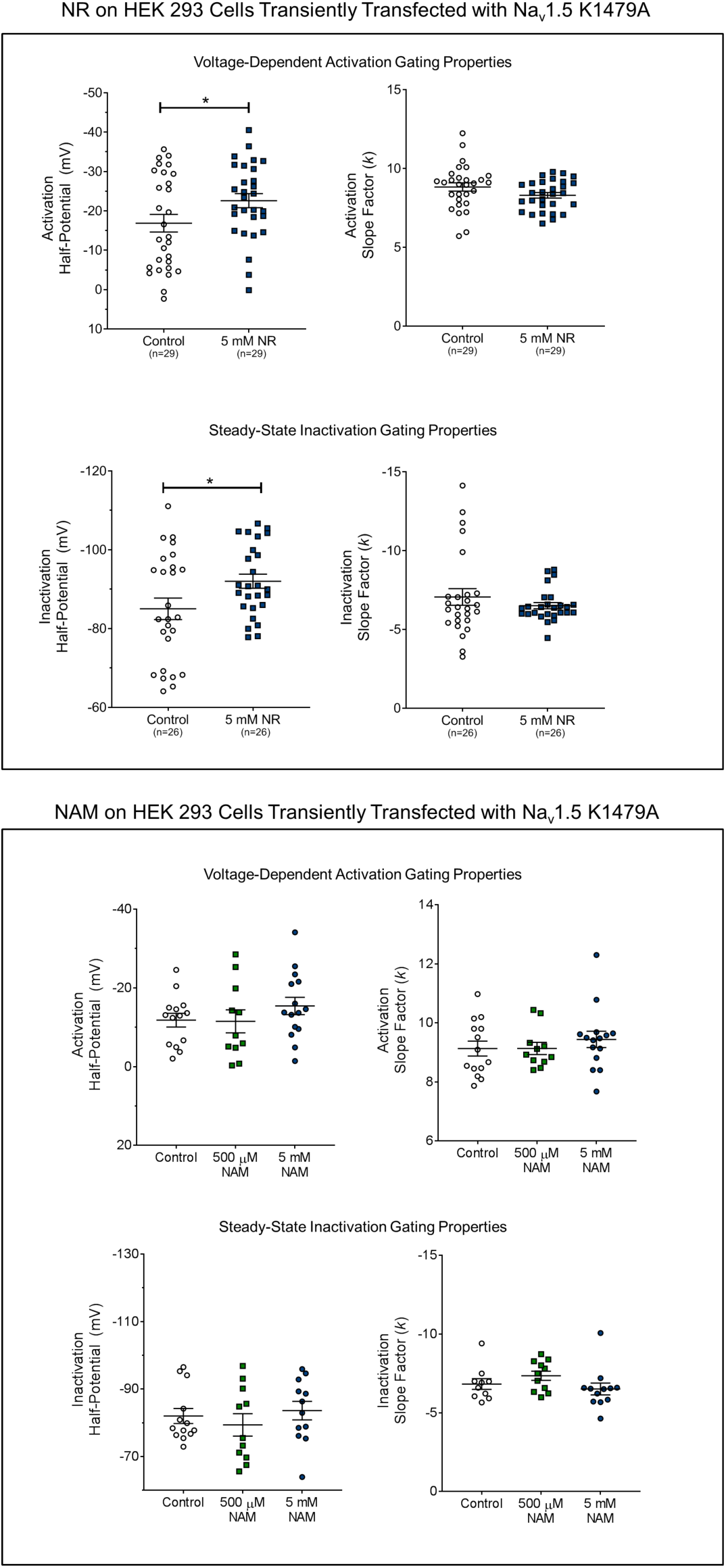
The Effect of Nicotinamide Riboside (NR) and Nicotinamide (NAM) on Na_V_1.5 K1479A Gating Properties in HEK293 Cells. Na_V_1.5 gating properties (including half-maximal potentials and slope factors) for steady-state activation and inactivation of HEK293 cells transiently transfected with Na_V_1.5 K1479A and treated with **(Top)** H_2_0 (vehicle) or 5 mM NR or **(Bottom)** H_2_O (vehicle), 500 µM NAM, or 5 mM NAM for 48 hours.

**Figure S13.**
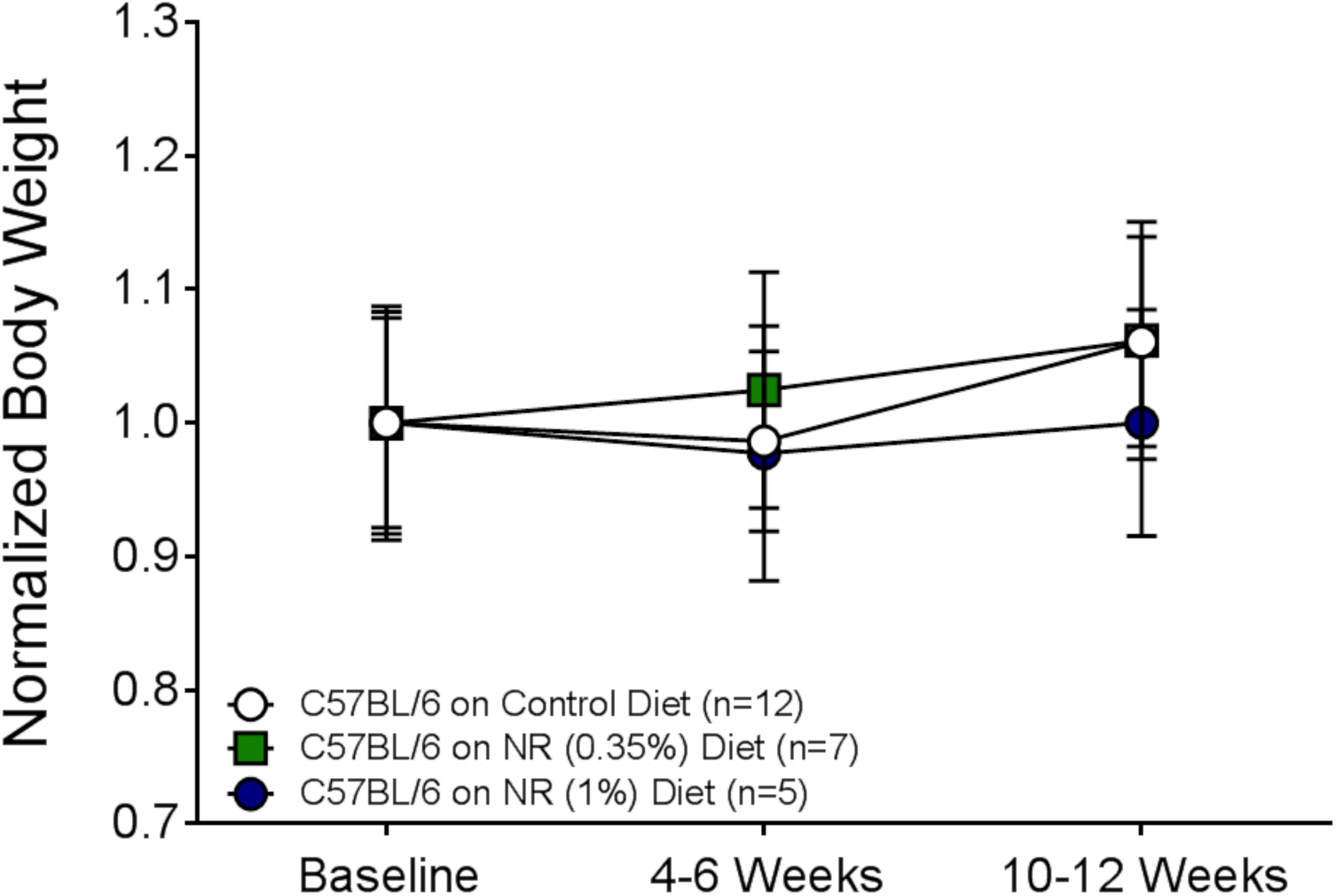
The Effect of Nicotinamide Riboside (NR) dietary supplementation on body weight in C57BL/6 wild-type mice. Body weights were assessed throughout the time course of NR supplementation. No changes in weight were observed between groups throughout the course of the experiment (Two-way ANOVA Bonferroni multiple comparisons post-hoc test).

**Table S1.**
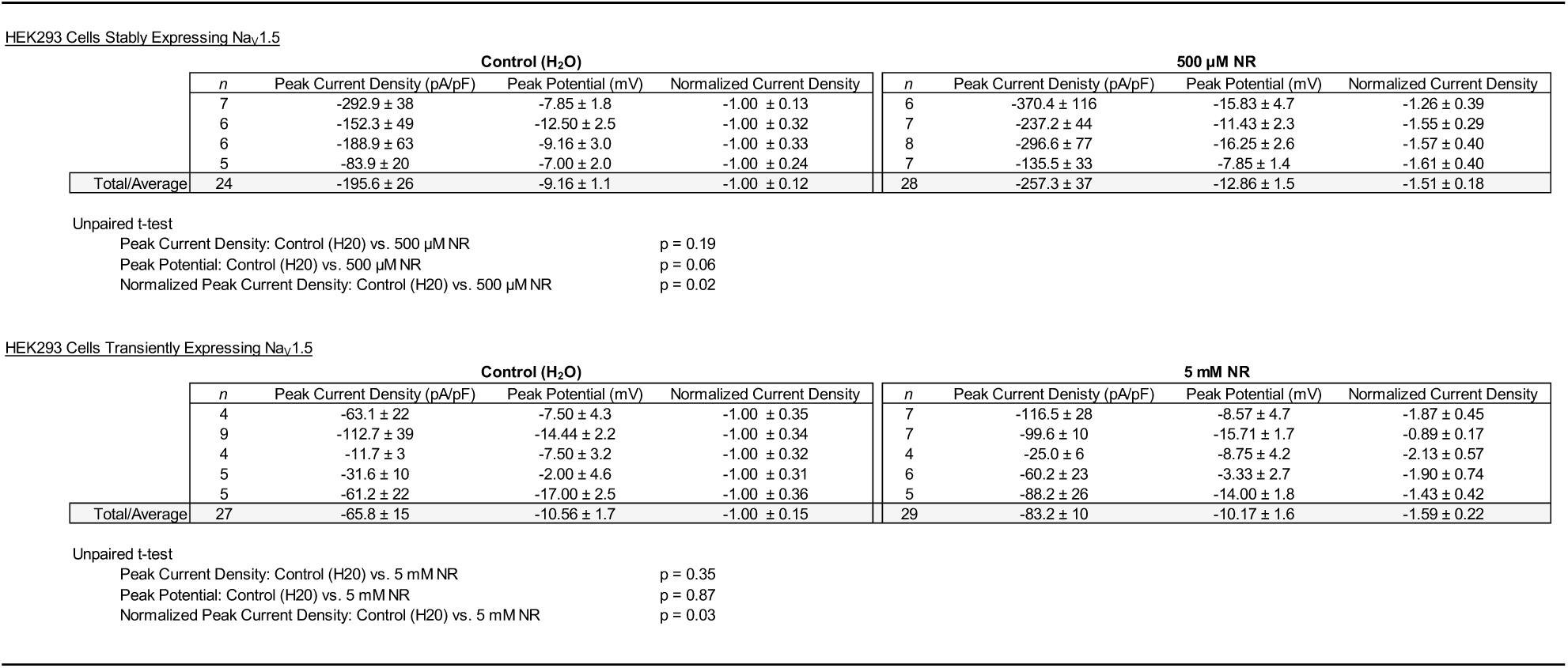
Current Densities, Peak Potentials, and Normalized Current Densities for HEK293 cells stably and transiently expressing Na_V_1.5 treated with Nicotinamide Riboside (NR).

**Table S2.**
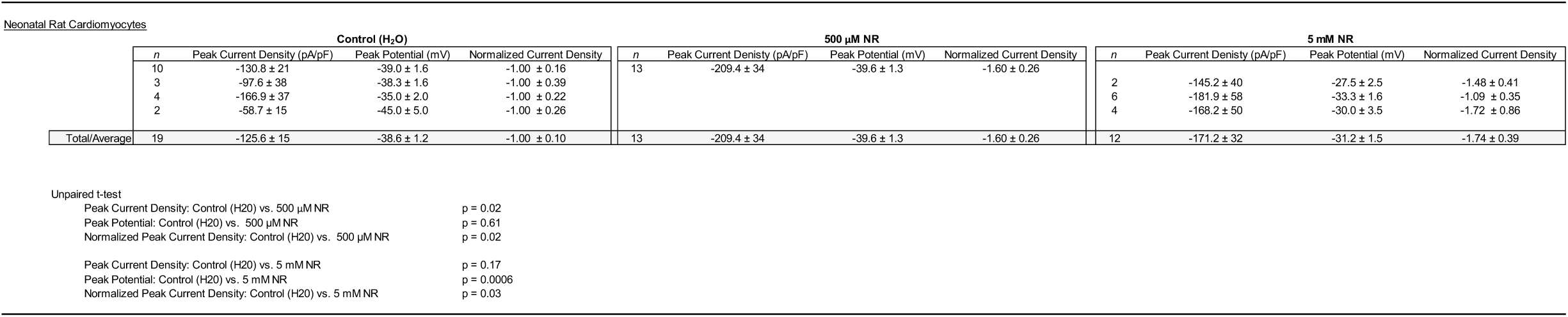
Current Densities, Peak Potentials, and Normalized Currents for Neonatal Rat Cardiomyocytes treated with Nicotinamide Riboside (NR).

**Table S3.**
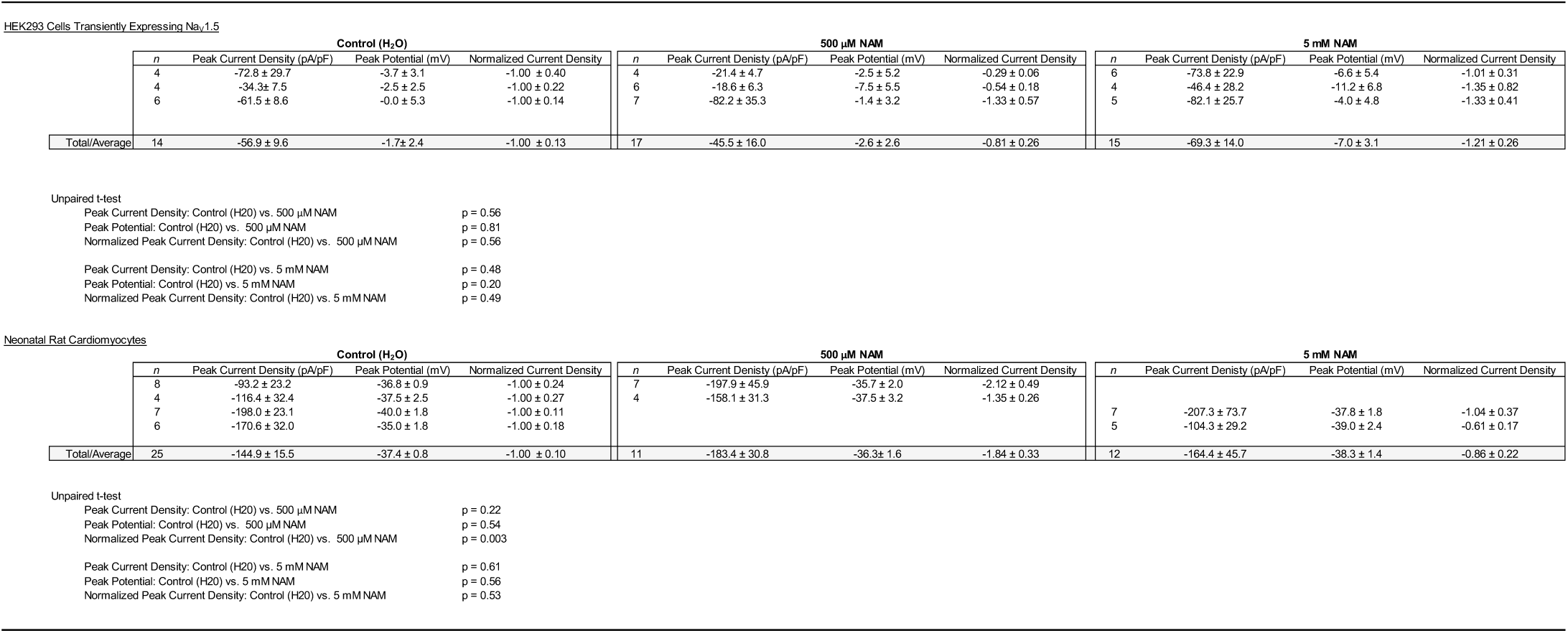
Current Densities, Peak Potentials, and Normalized Currents for HEK293 cells transiently expressing Na_V_1.5 and Neonatal Rat Cardiomyocytes treated with Nicotinamide (NAM).

**Table S4.**
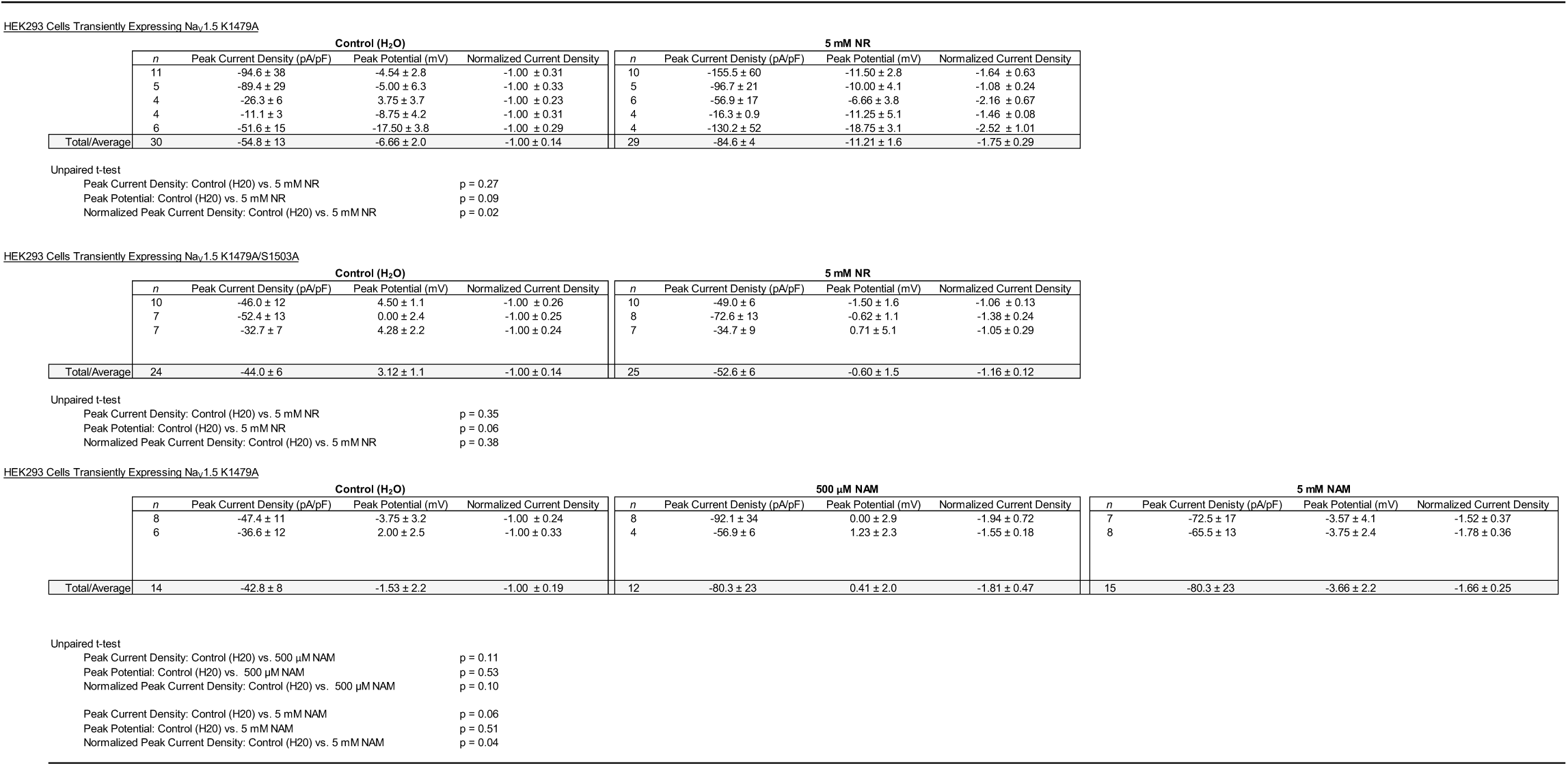
Current Densities, Peak Potentials, and Normalized Currents for HEK293 cells transiently expressing mutant forms (K1479A and K1479A/S1503A) and treated with either Nicotinamide Riboside (NR) or Nicotinamide (NAM).

**Table S5.** Gating Properties for Whole Cell Patch Clamp Experiments in HEK293 cells and Neonatal Rat Cardiomyocytes.

| HEK293 Cells Stably Expressing Nav1.5 |  |  | Activation Parameters |  | Inactivation Parameters |  |
| --- | --- | --- | --- | --- | --- | --- |
| Treatment Group | *n* | Peak $I_{Na}$ (Normalized Current) | Half Value (mV) | Slope (k) | Half Value (mV) | Slope (k) |
| Control (H<sub>2</sub>O) | 24 | -1.00 ± 0.12 | -20.04 ± 1.38 | 8.89 ± 0.35 | -78.44 ± 1.78 | -7.31 ± 0.23 |
| 500 μM NR | 28 | -1.51 ± 0.18 | -24.25 ± 1.33 | 8.73 ± 0.36 | -83.19 ± 2.22 | -7.10 ± 0.16 |
| HEK293 Cells Transiently Expressing Nav1.5 |  |  | Activation Parameters |  | Inactivation Parameters |  |
| Treatment Group | *n* | Peak $I_{Na}$ (Normalized Current) | Half Value (mV) | Slope (k) | Half Value (mV) | Slope (k) |
| Control (H<sub>2</sub>O) | 27 | -1.00 ± 0.15 | -22.15 ± 2.35 | 8.89 ± 0.35 | -83.66 ± 1.65 | -6.18 ± 0.22 |
| 5 mM NR | 29 | -1.59 ± 0.22 | -22.07 ± 1.64 | 8.73 ± 0.36 | -83.42 ± 1.47 | -6.39 ± 0.16 |
| HEK293 Cells Transiently Expressing Nav1.5 |  |  | Activation Parameters |  | Inactivation Parameters |  |
| Treatment Group | *n* | Peak $I_{Na}$ (Normalized Current) | Half Value (mV) | Slope (k) | Half Value (mV) | Slope (k) |
| Control (H<sub>2</sub>O) | 14 | -1.00 ± 0.13 | -12.23 ± 2.16 | 9.06 ± 0.26 | -77.72 ± 3.35 | -6.45 ± 0.40 |
| 500 μM NAM | 17 | -0.81 ± 0.26 | -12.80 ± 2.42 | 8.56 ± 0.22 | -74.51 ± 3.13 | -7.42 ± 0.42 |
| 5 mM NAM | 15 | -1.21 ± 0.26 | -18.42 ± 2.45 | 8.77 ± 0.34 | -80.78 ± 2.79 | -6.46 ± 0.37 |
| Rat Cardiomyocytes |  |  | Normalized Activation Parameters |  | Normalized Inactivation Parameters |  |
| Treatment Group | *n* | Peak $I_{Na}$ (Normalized Current) | Half Value (mV) | Slope | Half Value (mV) | Slope |
| Control (H<sub>2</sub>O) | 19 | -1.00 ± 0.10 | -49.17 ± 1.31 | 8.01 ± 0.38 | -97.49 ± 1.00 | -6.75 ± 0.33 |
| 500 μM NR | 13 | -1.60 ± 0.26 | -51.31 ± 1.16 | 6.93 ± 0.314 | -96.94 ± 0.83 | -7.31 ± 0.33 |
| 5 mM NR | 12 | -1.74 ± 0.39 | -42.36 ± 0.97 | 8.03 ± 0.18 | -92.91 ± 2.06 | -9.27 ± 0.29 |
| Treatment Group | *n* | Peak $I_{Na}$ (Normalized Current) | Half Value (mV) | Slope | Half Value (mV) | Slope |
| Control (H<sub>2</sub>O) | 25 | -1.00 ± 0.10 | -47.83 ± 1.47 | 8.62 ± 0.37 | -96.98 ± 1.37 | -6.50 ± 0.27 |
| 500 μM NAM | 11 | -1.84 ± 0.33 | -48.19 ± 1.92 | 7.93 ± 0.26 | -94.76 ± 2.43 | -6.91 ± 0.30 |
| 5 mM NAM | 12 | -0.86 ± 0.22 | -50.26 ± 2.74 | 7.05 ± 0.79 | -101.0 ± 2.60 | -6.05 ± 0.93 |
| Treatment Group | *n* | Peak $I_{Na}$ (Normalized Current) | Half Value (mV) | Slope | Half Value (mV) | Slope |
| Control (H<sub>2</sub>O) | 12 | -1.00 ± 0.11 | -49.21 ± 1.77 | 7.75 ± 0.18 | -95.15 ± 1.65 | -6.96 ± 0.24 |
| 500 μM meNAM | 6 | -0.93 ± 0.14 | -53.54 ± 2.36 | 7.75 ± 0.26 | -99.18 ± 2.38 | -6.56 ± 0.51 |
| 5 mM meNAM | 9 | -0.95 ± 0.09 | -45.19 ± 2.01 | 7.79 ± 0.21 | -91.47 ± 1.37 | -6.35 ± 0.15 |
| HEK293 Cells Transiently Expressing Nav1.5 K1479A |  |  |  |  |  |  |
| Treatment Group | *n* | Peak $I_{Na}$ (Normalized Current) | Half Value (mV) | Slope | Half Value (mV) | Slope |
| Control (H<sub>2</sub>O) | 29 | -1.00 ± 0.14 | -13.70 ± 2.38 | 8.72 ± 0.32 | -81.47 ± 2.73 | -7.51 ± 0.60 |
| 5 mM NR | 30 | -1.75 ± 0.29 | -21.29 ± 1.93 | 8.34 ± 0.18 | -90.42 ± 1.81 | -6.68 ± 0.20 |
| HEK293 Cells Transiently Expressing Nav1.5 K1479A/S1503A |  |  |  |  |  |  |
| Treatment Group | *n* | Peak $I_{Na}$ (Normalized Current) | Half Value (mV) | Slope | Half Value (mV) | Slope |
| Control (H<sub>2</sub>O) | 24 | -1.00 ± 0.14 | -7.89 ± 1.22 | 8.73 ± 0.19 | -75.43 ± 1.03 | -7.33 ± 0.27 |
| 5 mM NR | 25 | -1.16 ± 0.12 | -11.83 ± 1.45 | 9.07 ± 0.23 | -79.08 ± 1.21 | -7.39 ± 0.20 |
| HEK293 Cells Transiently Expressing Nav1.5 K1479A |  |  |  |  |  |  |
| Treatment Group | *n* | Peak $I_{Na}$ (Normalized Current) | Half Value (mV) | Slope | Half Value (mV) | Slope |
| Control (H<sub>2</sub>O) | 14 | -1.00 ± 0.19 | -11.81 ± 1.73 | 9.13 ± 0.25 | -82.03 ± 1.73 | -6.82 ± 0.33 |
| 500 μM NAM | 12 | -1.81 ± 0.47 | -11.51 ± 2.91 | 9.13 ± 0.20 | -79.39 ± 3.33 | -7.35 ± 0.29 |
| 5 mM NAM | 15 | -1.66 ± 0.25 | -15.41 ± 2.20 | 9.44 ± 0.27 | -83.63 ± 2.72 | -6.51 ± 0.37 |

**Table S6.** Effect of Nicotinamide Riboside on Cardiac Electrophysiology in C57BL/6 wild-type mice.

| **Baseline** |  |  |  |  |  |  |
| --- | --- | --- | --- | --- | --- | --- |
|  | Heart Rate<br>Lead I<br>(bpm) | Heart Rate<br>QT Measurement<br>(bpm) | PR Interval<br>(ms) | QRS Duration<br>(ms) | QT<br>(ms) | QTc<br>(ms) |
| Control Diet (n=12) | 530 ± 9 | 498 ± 11 | 46.7 ± 1.4 | 15.0 ± 0.2 | 94.2 ± 2.8 | 85.3 ± 2.1 |
| 0.35% NR Diet (n=7) | 539 ± 24 | 510 ± 21 | 49.1 ± 3.4 | 14.9 ± 0.5 | 103.0 ± 3.9 | 93.9 ± 2.3 |
| 1.0% NR Diet (n=5) | 552 ± 30 | 516 ± 18 | 46.8 ± 1.3 | 14.7 ± 0.4 | 88.1 ± 3.0 | 81.6 ± 2.8 |
| **5-6 Weeks Post-Diet** |  |  |  |  |  |  |
|  | Lead I Heart Rate<br>(bpm) | Heart Rate<br>(bpm) | PR Interval<br>(ms) | QRS Duration<br>(ms) | QT<br>(ms) | QTc<br>(ms) |
| Control Diet (n=12) | 534 ± 15 | 492 ± 10 | 47.6 ± 1.9 | 14.7 ± 0.3 | 92.7 ± 2.0 | 83.9 ± 2.0 |
| 0.35% NR Diet (n=7) | 537 ± 14 | 502 ± 11 | 45.8 ± 2.2 | 14.5 ± 0.5 | 99.1 ± 2.3 | 90.4 ± 1.8 |
| 1.0% NR Diet (n=5) | 566 ± 21 | 535 ± 32 | 46.6 ± 2.7 | 13.7 ± 0.2 | 85.6 ± 3.9 | 79.8 ± 2.1 |
| **10-12 Weeks Post-Diet** |  |  |  |  |  |  |
|  | Lead I Heart Rate<br>(bpm) | Heart Rate<br>(bpm) | PR Interval<br>(ms) | QRS Duration<br>(ms) | QT<br>(ms) | QTc<br>(ms) |
| Control Diet (n=12) | 532 ± 12 | 516 ± 11 | 45.9 ± 1.6 | 14.5 ± 0.3 | 94.0 ± 2.4 | 86.9 ± 1.9 |
| 0.35% NR Diet (n=7) | 558 ± 16 | 526 ± 11 | 48.0 ± 2.6 | 15.2 ± 0.4 | 95.6 ± 2.4 | 89.3 ± 1.8 |
| 1.0% NR Diet (n=5) | 502 ± 35 | 476 ± 24 | 42.3 ± 1.1 | 14.8 ± 0.4 | 83.4 ± 4.0 | 73.8 ± 2.2 |

**Table S7.** Effect of Nicotinamide Riboside on Cardiac Function in C57BL/6 wild-type mice. Heart Rate, End-diastolic volume (EDV), End-systolic volume (ESV), Mass, Stroke volume (SV), Cardiac Output (CO), Ejection Fraction (EF).

|  | Baseline |  |  |  |  |  |  |
| --- | --- | --- | --- | --- | --- | --- | --- |
| | Heart Rate<br>(beats/min) | EDV<br>(μL) | ESV<br>(μL) | Mass<br>(mg) | SV<br>(μL) | CO<br>(μL/min) | EF<br>(%) |
| Control Diet (n=12) | 679 ± 9. | 27.4 ± 2.1 | 5.3 ± 0.5 | 74.9 ± 4.4 | 22.1 ± 2.2 | 14888 ± 1399 | 79 ± 3 |
| 0.35% NR Diet (n=7) | 701 ± 12 | 23.7 ± 3.7 | 4.3 ± 0.6 | 70.6 ± 2.9 | 18.3 ± 3.9 | 12802 ± 2727 | 73 ± 6 |
| 1.0% NR Diet (n=5) | 686 ± 19 | 32.5 ± 3.7 | 5.3 ± 0.6 | 82.1 ± 5.4 | 28.2 ± 2.9 | 19352 ± 2041 | 87 ± 2 |
|  | 10-12 Weeks |  |  |  |  |  |  |
| | Heart Rate<br>(beats/min) | EDV<br>(μL) | ESV<br>(μL) | Mass<br>(mg) | SV<br>(μL) | CO<br>(μL/min) | EF<br>(%) |
| Control Diet (n=12) | 691 ± 10 | 25.4 ± 1.5 | 4.9 ± 0.5 | 72.5 ± 3.4 | 20.4 ± 1.3 | 14095 ± 928 | 80 ± 2 |
| 0.35% NR Diet (n=7) | 637 ± 46 | 24.8 ± 2.4 | 4.9 ± 0.7 | 72.5 ± 3.1 | 19.8 ± 2.3 | 12259 ± 1418 | 79 ± 3 |
| 1.0% NR Diet (n=5) | 675 ± 16 | 28.4 ± 5.9 | 5.2 ± 1.3 | 77.0 ± 9.1 | 19.8 ± 2.3 | 15699 ± 3276 | 82 ± 2 |

## REFERENCES

1. Veerman CC, Wilde AA and Lodder EM. The cardiac sodium channel gene SCN5A and its gene product NaV1.5: Role in physiology and pathophysiology. Gene. 2015;573:177–87.

2. Brugada J and Brugada P. Further characterization of the syndrome of right bundle branch block, ST segment elevation, and sudden cardiac death. J Cardiovasc Electrophysiol. 1997;8:325–31.

3. Grant AO. Electrophysiological basis and genetics of Brugada syndrome. J Cardiovasc Electrophysiol. 2005;16 Suppl 1:S3–7.

4. Wang DW, Viswanathan PC, Balser JR, George AL, Jr. and Benson DW. Clinical, genetic, and biophysical characterization of SCN5A mutations associated with atrioventricular conduction block. Circulation. 2002;105:341–6.

5. Yan GX and Antzelevitch C. Cellular basis for the Brugada syndrome and other mechanisms of arrhythmogenesis associated with ST-segment elevation. Circulation. 1999;100:1660–6.

6. Liu M, Yang KC and Dudley SC, Jr. Cardiac Sodium Channel Mutations: Why so Many Phenotypes? Curr Top Membr. 2016;78:513–59.

7. Remme CA. Cardiac sodium channelopathy associated with SCN5A mutations: electrophysiological, molecular and genetic aspects. J Physiol. 2013;591:4099–116.

8. Savio-Galimberti E, Argenziano M and Antzelevitch C. Cardiac Arrhythmias Related to Sodium Channel Dysfunction. Handb Exp Pharmacol. 2018;246:331–354.

9. Makielski JC. Late sodium current: A mechanism for angina, heart failure, and arrhythmia. Trends Cardiovasc Med. 2016;26:115–22.

10. Remme CA and Bezzina CR. Sodium channel (dys)function and cardiac arrhythmias. Cardiovasc Ther. 2010;28:287–94.

11. Matasic DS, Brenner C and London B. Emerging potential benefits of modulating NAD(+) metabolism in cardiovascular disease. Am J Physiol Heart Circ Physiol. 2018;314:H839–H852.

12. Liu M, Sanyal S, Gao G, Gurung IS, Zhu X, Gaconnet G, Kerchner LJ, Shang LL, Huang CL, Grace A, London B and Dudley SC, Jr. Cardiac Na+ current regulation by pyridine nucleotides. Circ Res. 2009;105:737–45.

13. Liu M, Shi G, Yang KC, Gu L, Kanthasamy AG, Anantharam V and Dudley SC, Jr. Role of protein kinase C in metabolic regulation of the cardiac Na(+) channel. Heart Rhythm. 2017;14:440–447.

14. Valdivia CR, Ueda K, Ackerman MJ and Makielski JC. GPD1L links redox state to cardiac excitability by PKC-dependent phosphorylation of the sodium channel SCN5A. Am J Physiol Heart Circ Physiol. 2009;297:H1446–52.

15. Vikram A, Lewarchik CM, Yoon JY, Naqvi A, Kumar S, Morgan GM, Jacobs JS, Li Q, Kim YR, Kassan M, Liu J, Gabani M, Kumar A, Mehdi H, Zhu X, Guan X, Kutschke W, Zhang X, Boudreau RL, Dai S, Matasic DS, Jung SB, Margulies KB, Kumar V, Bachschmid MM, London B and Irani K. Sirtuin 1 regulates cardiac electrical activity by deacetylating the cardiac sodium channel. Nat Med. 2017;23:361–367.

16. Xu Q, Patel D, Zhang X and Veenstra RD. Changes in cardiac Nav1.5 expression, function, and acetylation by pan-histone deacetylase inhibitors. Am J Physiol Heart Circ Physiol. 2016;311:H1139–H1149.

17. Nicotinamide Riboside in LVAD Recipients (PilotNR-LVAD, NCT03727646). US National Library of Medicine. 2018;clinicaltrials.gov.

18. Nicotinamide Riboside in Systolic Heart Failure (NCT03423342). US National Library of Medicine. 2018;clinicaltrials.gov.

19. Effects of Nicotinamide Riboside on Metabolism and Vascular Function (NCT03501433). US National Library of Medicine. 2018;clinicaltrials.gov.

20. Nicotinamide Riboside for Treating Elevated Systolic Blood Pressure and Arterial Stiffness in Middle-aged and Older Adults (NCT03821623). US National Library of Medicine. 2019;clinicaltrials.gov.

21. Trammell SA and Brenner C. Targeted, LCMS-based Metabolomics for Quantitative Measurement of NAD(+) Metabolites. Comput Struct Biotechnol J. 2013;4:e201301012.

22. London B, Jeron A, Zhou J, Buckett P, Han X, Mitchell GF and Koren G. Long QT and ventricular arrhythmias in transgenic mice expressing the N terminus and first transmembrane segment of a voltage-gated potassium channel. Proc Natl Acad Sci U S A. 1998;95:2926–31.

23. Mitchell GF, Jeron A and Koren G. Measurement of heart rate and Q-T interval in the conscious mouse. Am J Physiol. 1998;274:H747–51.

24. Rasmussen TP, Wu Y, Joiner ML, Koval OM, Wilson NR, Luczak ED, Wang Q, Chen B, Gao Z, Zhu Z, Wagner BA, Soto J, McCormick ML, Kutschke W, Weiss RM, Yu L, Boudreau RL, Abel ED, Zhan F, Spitz DR, Buettner GR, Song LS, Zingman LV and Anderson ME. Inhibition of MCU forces extramitochondrial adaptations governing physiological and pathological stress responses in heart. Proc Natl Acad Sci U S A. 2015;112:9129–34.

25. Liu M, Gu L, Sulkin MS, Liu H, Jeong EM, Greener I, Xie A, Efimov IR and Dudley SC, Jr. Mitochondrial dysfunction causing cardiac sodium channel downregulation in cardiomyopathy. J Mol Cell Cardiol. 2013;54:25–34.

26. Ratajczak J, Joffraud M, Trammell SA, Ras R, Canela N, Boutant M, Kulkarni SS, Rodrigues M, Redpath P, Migaud ME, Auwerx J, Yanes O, Brenner C and Canto C. NRK1 controls nicotinamide mononucleotide and nicotinamide riboside metabolism in mammalian cells. Nat Commun. 2016;7:13103.

27. Herfst LJ, Rook MB and Jongsma HJ. Trafficking and functional expression of cardiac Na+ channels. J Mol Cell Cardiol. 2004;36:185–93.

28. Zhang X, Yoon JY, Morley M, McLendon JM, Mapuskar KA, Gutmann R, Mehdi H, Bloom HL, Dudley SC, Ellinor PT, Shalaby AA, Weiss R, Tang WHW, Moravec CS, Singh M, Taylor AL, Yancy CW, Feldman AM, McNamara DM, Irani K, Spitz DR, Breheny P, Margulies KB, London B and Boudreau RL. A common variant alters SCN5A-miR-24 interaction and associates with heart failure mortality. J Clin Invest. 2018;128:1154–1163.

29. Aiba T, Farinelli F, Kostecki G, Hesketh GG, Edwards D, Biswas S, Tung L and Tomaselli GF. A mutation causing Brugada syndrome identifies a mechanism for altered autonomic and oxidant regulation of cardiac sodium currents. Circ Cardiovasc Genet. 2014;7:249–56.

30. Zhou J, Shin HG, Yi J, Shen W, Williams CP and Murray KT. Phosphorylation and putative ER retention signals are required for protein kinase A-mediated potentiation of cardiac sodium current. Circ Res. 2002;91:540–6.

31. Tateyama M, Rivolta I, Clancy CE and Kass RS. Modulation of cardiac sodium channel gating by protein kinase A can be altered by disease-linked mutation. J Biol Chem. 2003;278:46718–26.

32. Bai P. Biology of Poly(ADP-Ribose) Polymerases: The Factotums of Cell Maintenance. Mol Cell. 2015;58:947–58.

33. Virag L and Szabo C. The therapeutic potential of poly(ADP-ribose) polymerase inhibitors. Pharmacol Rev. 2002;54:375–429.

34. Camacho-Pereira J, Tarrago MG, Chini CCS, Nin V, Escande C, Warner GM, Puranik AS, Schoon RA, Reid JM, Galina A and Chini EN. CD38 Dictates Age-Related NAD Decline and Mitochondrial Dysfunction through an SIRT3-Dependent Mechanism. Cell Metab. 2016;23:1127–1139.

35. Kulikova V, Shabalin K, Nerinovski K, Yakimov A, Svetlova M, Solovjeva L, Kropotov A, Khodorkovskiy M, Migaud ME, Ziegler M and Nikiforov A. Degradation of Extracellular NAD(+) Intermediates in Cultures of Human HEK293 Cells. Metabolites. 2019;9.

36. Moreno JD and Clancy CE. Pathophysiology of the cardiac late Na current and its potential as a drug target. J Mol Cell Cardiol. 2012;52:608–19.

37. Rivaud MR, Baartscheer A, Verkerk AO, Beekman L, Rajamani S, Belardinelli L, Bezzina CR and Remme CA. Enhanced late sodium current underlies pro-arrhythmic intracellular sodium and calcium dysregulation in murine sodium channelopathy. Int J Cardiol. 2018;263:54–62.

38. Song Y, Shryock JC and Belardinelli L. An increase of late sodium current induces delayed afterdepolarizations and sustained triggered activity in atrial myocytes. Am J Physiol Heart Circ Physiol. 2008;294:H2031–9.

39. Nagatomo T, January CT and Makielski JC. Preferential block of late sodium current in the LQT3 DeltaKPQ mutant by the class I(C) antiarrhythmic flecainide. Mol Pharmacol. 2000;57:101–7.

40. Doshi D and Morrow JP. Potential application of late sodium current blockade in the treatment of heart failure and atrial fibrillation. Rev Cardiovasc Med. 2009;10 Suppl 1:S46–52.

41. Fukaya H, Plummer BN, Piktel JS, Wan X, Rosenbaum DS, Laurita KR and Wilson LD. Arrhythmogenic cardiac alternans in heart failure is suppressed by late sodium current blockade by ranolazine. Heart Rhythm. 2019;16:281–289.

42. Valdivia CR, Chu WW, Pu J, Foell JD, Haworth RA, Wolff MR, Kamp TJ and Makielski JC. Increased late sodium current in myocytes from a canine heart failure model and from failing human heart. J Mol Cell Cardiol. 2005;38:475–83.

43. Liu M, Liu H and Dudley SC, Jr. Reactive oxygen species originating from mitochondria regulate the cardiac sodium channel. Circ Res. 2010;107:967–74.

44. Xie LH, Chen F, Karagueuzian HS and Weiss JN. Oxidative-stress-induced afterdepolarizations and calmodulin kinase II signaling. Circ Res. 2009;104:79–86.

45. Kilfoil PJ, Tipparaju SM, Barski OA and Bhatnagar A. Regulation of ion channels by pyridine nucleotides. Circ Res. 2013;112:721–41.

46. Salama G and London B. Mouse models of long QT syndrome. J Physiol. 2007;578:43–53.

